# Silencing ABAP1 INTERACTING PROTEIN 10 (AIP10) enhances root colonization by beneficial bacteria and improves plant performance under nutrient-limited conditions in *Arabidopsis thaliana*

**DOI:** 10.64898/2026.01.25.701605

**Authors:** Maria Clara Urquiaga, Helkin Giovani Forero Ballesteros, João Victor Silva, Patrícia Montessoro, Sara Gawantka Evangelista, Adriana Silva Hemerly

## Abstract

Sustaining high agricultural productivity with minimal environmental impact requires innovative and sustainable strategies that reduce reliance on mineral fertilizers. Promoting root association with plant growth-promoting bacteria (PGPB), either within the native microbiome or through bioinoculant application, represents a promising strategy to improve crop performance while reducing mineral fertilizer inputs. The success of this strategy, however, is strongly influenced by plant genetic traits that regulate microbial recruitment and colonization. Here, we tested whether silencing the ABAP1 Interacting Protein (AIP10), a negative regulator that links cell division with primary metabolism, modulates the association of *Arabidopsis thaliana* to PGPB. Non-inoculated *aip10-1* roots exhibited gene expression patterns similar to genotypes with enhanced microbial associations. *AIP10* silencing reshaped root and rhizosphere bacterial communities, favoring beneficial PGPB associations and limiting potential pathogens. Consistently, *aip10-1* plants showed greater colonization by inoculated diazotrophic PGPB, particularly in low fertilization conditions, leading to increased plant performance. These effects were accompanied by modulation of plant cell cycle and nitrogen assimilation pathways, together with increased bacterial colonization and *nifH* expression. Our findings suggest that AIP10 functions as a regulatory hub coordinating growth and metabolism with beneficial PGPB recruitment. Modulating AIP10 could enhance plant productivity and support more sustainable and regenerative agriculture practices.

**Highlight:** AIP10 silencing participates in a regulatory hub coordinating plant cell cycle and metabolism with recruitment of beneficial bacteria in the root microbiota, contributing to improved plant growth and productivity under nutrient-limited conditions.

## 1. Introduction

Global population growth has increased the demand for agricultural products, including food, bioenergy, and other bioproducts, while climate change negatively impacts agricultural productivity. To sustain high yields while protecting the environment, modern agriculture must develop innovative and sustainable technologies that reduce environmental impacts, particularly those associated with the excessive use of chemical fertilizers, which contribute to increased greenhouse gas (GHG) emissions ((Tilman *et al*., 2011). In parallel, nitrogen use efficiency (NUE) is estimated to remain below 50% for many crop species ((Mueller *et al*., 2019; Govindasamy *et al*., 2023). Bioinoculants based on non-nodulating Plant Growth Promoting Bacteria (PGPB) represent a sustainable alternative to mineral fertilization, as they increase nutrients availability, improving soil fertility mainly through biological nitrogen fixation (BNF), mineral solubilization and iron-chelating siderophores production (Carvalho *et al*., 2016; Omara *et al*., 2019; Zuluaga *et al*., 2020; Favero *et al*., 2021; Pereira *et al*., 2021; Rodrigues *et al*., 2022) Furthermore, PGPB can stimulate plant development by synthesizing phytohormones, modulating antioxidant production, and altering the expression of defense-related genes, thereby enhancing plant tolerance to both biotic and abiotic stresses (Vargas *et al*., 2014; Fukami *et al*., 2017; Filgueiras *et al*., 2020; Moretti *et al*., 2021; Nunes *et al*., 2021; Orozco-Mosqueda *et al*., 2023).

Agriculturally important non-nodulating monocot crops, such as maize, rice and sugarcane, have shown beneficial responses to inoculation with bioinoculants based on non-nodulating diazotrophic bacteria (Hardoim *et al*., 2020; Pankievicz *et al*., 2021; Carvalho *et al*., 2022; Ferrarezi *et al*., 2022; Tian *et al*., 2022). Although BNF in these crops is less efficient than in legumes, numerous studies have demonstrated that bacterial inoculation significantly increases grain yield and biomass with BNF contributing up to 37% of the total N in maize (Alves *et al*., 2014; Di Salvo *et al*., 2018; dos Santos *et al*., 2020; Aquino *et al*., 2021; Dias *et al*., 2021). The use of PGPB bioinoculants represents a promising strategy for developing more resilient and productive crops. However, their efficient application still faces major challenges. The success of these interactions depends on both biotic and abiotic environmental factors, as well as on the host plant genotype by modulation of plant gene expression (Carvalho *et al*., 2011, 2014; Vargas *et al*., 2012). This modulation allows the host plant to recognize the bacterial strain as a beneficial partner and, in turn, regulate plant gene expression to facilitate PGPB colonization, thereby directly or indirectly activating plant pathways that promote plant growth (Carvalho *et al*., 2011; Do Amaral *et al*., 2014; Espindula *et al*., 2019; Hardoim *et al*., 2020; Ballesteros *et al*., 2021; Pankievicz *et al*., 2021; Thiebaut *et al*., 2022; Rosman *et al*., 2024).

Alterations in plant genes were shown to influence the assembly of root-associated bacterial communities, indicating that root microbiome composition is strongly shaped by plant genotype (Zang *et al.,* 2023). Modulation of defense-related pathways, such as salicylic acid signaling, has been reported to affect the recruitment of specific bacterial groups in the rhizosphere (Lebeis *et al*., 2015). Likewise, genes involved in key signaling modules including the PHR1–RALF–FERONIA pathway have been implicated in shaping root-associated microbial communities with distinct functional capacities (Tang *et al*., 2022). Additionally, genes regulating the production of root-derived metabolites contribute to differences in microbial composition and the enrichment of plant growth–promoting taxa (Stringlis *et al*., 2018). These studies highlight that genetic variation in plants, including reductions in gene activity, can play a significant role in determining the structure and function of the root microbiome.

Plant plasticity, which allows the modulation of plant growth in response to different environmental cues, including association with PGPB that enhance plant biomass and productivity. Ultimately, biomass increases are driven by higher rates the regulation of cell division rates (Qi and Zhang, 2020; Pierik *et al*., 2021; Zluhan-Martínez *et al*., 2021). Recently, our group described a novel cell cycle regulator, the ABAP1 Interacting Protein 10 (AIP10), which acts in a central hub integrating cell division rates with transcriptional reprogramming of basal metabolism (Montessoro *et al*., 2025). This role is mediated by AIP10 protein interactions with the Armadillo BTB Arabidopsis protein 1 (ABAP1), a negative regulator of the cell cycle in plants, with KIN10, a catalytic subunit of the Sucrose non-fermenting-1-related protein kinase 1 (SnRK1), and possibly additional partners (Masuda *et al*., 2008; Montessoro *et al*., 2025). In *Arabidopsis thaliana, aip10-1* knockout mutant plants exhibited increased cell division rates, resulting in enhanced shoot and root vegetative growth, as well as higher numbers of silique and seed production (Montessoro *et al*., 2025). In addition, reduced *AIP10* expression triggered a major transcriptional reprogramming of plant primary metabolism mediated by SnRK1, leading to increased photosynthetic and water use efficiency and greater carbon fixation that met plant energy demands for increased biomass accumulation, and higher contents of proteins, lipids and carbohydrates (Montessoro *et al*., 2025).

Developing strategies that enhance plant-bacteria interaction is a promising approach to increasing agricultural productivity and promoting more sustainable agriculture. *A. thaliana* was already used as a model system to assess developmental responses and proteomic changes triggered by association with PGPB, such as endophytic diazotrophic bacteria (Cohen *et al*., 2015; Souza *et al*., 2016; Leandro *et al*., 2019; Dos Santos *et al*., 2020). Here, we evaluated the role of AIP10 in interactions with PGPB, both within the native root microbiota and following inoculation with PGPB-based inoculants. The previously described *aip10-1* full knockout mutant was used as a functional model and investigated using multiple complementary approaches, including RT-qPCR and RNA-seq gene expression profiling, root and soil microbiome metabarcoding, PGPB inoculation and comprehensive plant phenotyping. The transcriptional profile of non-inoculated *aip10-1* roots resembled signatures of genotypes with enhanced capacity to interact with beneficial microorganisms. Loss of *AIP10* function reshaped root-associated bacterial communities, in roots and rhizospheric soil, favoring the recruitment of beneficial PGPB species while reducing potentially pathogenic taxa. Accordingly, *aip10-1* plants showed higher colonization by inoculated diazotrophic PGPB, leading to improved vegetative and reproductive growth. Moreover, under low fertilizer conditions, *aip10-1* plants exhibited a stronger response to PGPB bioinoculants, achieving biomass levels similar to non-inoculated plants grown under high fertilization. These findings indicate that AIP10 silencing is a promising strategy to enhance PGPB colonization and improve nutrient use efficiency under reduced chemical fertilization, contributing to a more sustainable and regenerative agriculture.

## 2. Material and Methods

### 2.1. Bacterial strains and growth conditions

For plant inoculation, the diazotrophic bacterial strain *Gluconacetobacter diazotrophicus* PAL5 DsRed, labeled with red fluorescent protein and *Herbaspirillum seropedicae* RAM10, labeled with green fluorescent protein, kindly provided by Dr. Gonçalo Apolinário de Souza Filho (State University of Norte Fluminense Darcy Ribeiro), and *Herbaspirillum seropedicae* HRC54 (BR11335), obtained from EMBRAPA Agrobiologia (Rio de Janeiro, Brazil), were used. Additionally, the commercial biofertilizer BioFree® (liquid formulation), supplied by BioFROST and registered with the Brazilian Ministry of Agriculture, Livestock and Food Supply (MAPA, No. PR001593-8.000001), containing *Azospirillum brasilense* Ab-V6 and *Pseudomonas fluorescens* CCTB03 (1×10¹¹ CFU/L), was employed in the assays. *G. diazotrophicus* PAL5 DsRed was cultured in LGI-P medium (100 g/L granulated sugar; 2 mL/L K_2_HPO sol. 10%; 6mL/L KH_2_PO_4_ sol. 10%; 2 mL/L MgSO_4_.7H_2_O sol. 10%; 2 mL/L CaCl_2_.2H_2_O sol. 1%; 1 ml/L FeCl_3_.6H_2_O sol. 1%; 2 mL/L Na_2_MoO_4_.2H_2_O sol. 0.1%; 5 mL/L bromothymol blue sol. 5% w/v in 0.2N KOH; pH 5.8) supplemented with streptomycin (50 mg/L) and 1 mL of vitamin stock solution (biotin 100mg/L; pyridoxal-HCL 200 mg/L) per liter of medium (Baldani *et al*., 1996) and *H. seropedicae* HRC54 was grown in JNFB medium (5 g/L malic acid; 6 mL/L K_2_HPO sol. 10%; 18 mL/L KH_2_PO_4_ sol. 10%; 2 mL/L MgSO_4_.7H_2_O sol. 10%; 1 mL/L NaCl sol. 10%; 2 mL/L bromothymol blue sol. 5% w/v in 0.2N KOH; 4 mL/L FeEDTA sol. 1.64%; 4 g/L KOH; 1,07 g/L NH_4_Cl; pH 6.0) supplemented with 1 mL of vitamin stock solution and 2 mL of micronutrients stock solution (1 g/L Na_2_MoO_4_.2H_2_O; 1.175 g/L MnSO.HO; 1.4 g/L HBO; 0.04 g/L CuSO.5HO; 0.12 g/L ZnSO.7HO) per liter (Baldani *et al*., 1996). Both strains were subsequently transferred to DYGS liquid medium (2 g/L malic acid; 2 g/L glucose; 1.5 g/L bacteriological peptone; 2 g/L yeast extract; 0.5 g/L K_2_HPO_4_; 0.5 g/L MgSO_4_.7H_2_O and 1.5 g/L glutamic acid, pH 6.0 for HRC54 and pH 5.8 for Pal5 without malic acid) (Baldani *et al*., 1996) and incubated at 30 °C with agitation (200 rpm) for 6–12 hours until reaching an optical density at 660 nm of approximately 0.8, corresponding to ∼10⁸ cells mL^-1^. For plant inoculation, bacterial cultures and the commercial inoculant were diluted 1:1000 (v/v) in autoclaved distilled water.

### 2.2. Plant material and inoculation with plant-growth promoting bacteria

Seeds of the wild-type *A. thaliana* ecotype Col-0 and the knockout mutant line *aip10-1* (SALK_022332, T-DNA insertion), obtained from the Salk collection (Salk Institute Genomic Analysis Laboratory) and previously characterized (Montessoro *et al*., 2025), were used for the AIP10 functional analyses. The seeds were sterilized by exposure to chlorine gas, generated by the reaction between 97 mL of bleach (4.3%) and 3 mL of hydrochloric acid (37%). After 2 hrs of treatment, the seeds were left in a laminar flow hood until the gas had completely evaporated. Then, they were sown in Petri dishes containing Murashige and Skoog (MS) ½ medium, supplemented with 0.05% 2-(N-morpholino) ethanesulfonic acid (MES), 1% sucrose and 0.8% Phytagel (Sigma-Aldrich®), adjusted to pH 5.7. Seeds were stratified at 4 °C for 2 days and subsequently grown vertically in a growth chamber with a 12/12 hrs photoperiod at 21 °C.

Four days after germination (4 DAG), 2 μL of the diluted bacterial suspension (1:1000), containing approximately 10² cells μL^-1^, were applied to the root zone. Control seedlings were mock-treated with sterile DYGS medium diluted in the same proportion. After inoculation, the seedlings were transferred to Petri dishes containing modified MS ½ medium (sucrose-free) and maintained under the same culture conditions for another 7 days, totaling 11 DAG. For molecular analyses, three biological replicates per treatment were harvested at 11 DAG, with shoots and roots separated, frozen in liquid nitrogen, and stored at −80 °C. Plants were then transplanted into pots containing a standard mixture of substate:vermiculite (3:1, w/w) and grown under a 16/8 hrs photoperiod at 21 °C until 17 DAG.

In the experiment with different nutritional doses, after a 6-day acclimatization period, the plants were watered with two different concentrations of NPK 20-20-20 fertilizer, defined as low nutrient availability (final concentration of 0.06 g/L, corresponding to 12 mg/L N, 5.2 mg/L P and 10 mg/L K) and high nutrient availability (final concentration of 0.12 g/L, corresponding to 24 mg/L N, 10.4 mg/L P and 20 mg/L K).The plants were harvested 31 days after inoculation (31 DAI), corresponding to 35 DAG, for phenotypic analysis, and their productivity was evaluated at 60 DAG.

### 2.3. Phenotypic analyses of A. thaliana plants

Phenotypic analyses were performed on *A. thaliana* plants after separating shoots and roots at 31 DAI. Fresh matter weight was measured using an analytical precision balance with eight biological replicates. Thereafter, shoot and root samples were then placed in paper bags and oven-dried at 60 °C for 6 days. Dry weight was subsequently measured using the same balance, with eight biological replicates, to determine biomass. In addition, the rosette area was determined from images evaluated in imageJ software (Schindelin *et al*., 2012). The values of fresh and dry matter and rosette area obtained were statistically analyzed using GraphPad Prism software version 8.0.0. Statistical analysis was performed using Student’s T-test. Statistical significance was defined in all cases as *p*-value < 0.05

To analyze root development, harvested roots were immersed in 70% ethanol for 7 days. Subsequently, roots were laid out in an acrylic container (10 × 45 cm), with water at an approximate depth of 1 cm for scanning. They were then characterized by image analysis using WinRHIZO Pro® software (Regent Instruments, QC, Quebec, Canada) coupled to an Epson Expression 11000XL LA2400 image scanner, as described in previous works (Bauhus and Messier, 1999). Root length (cm) and surface area (m²) were evaluated in five plants per treatment. All analyzes used the Student’s T-test. Statistical significance was defined as *p*-value < 0.05.

Productivity analyses were performed at 56 DAI (60 DAG), including measurements of number of siliques, length of the main inflorescence axis, and weight of seeds per plant. The resulting data were subjected to statistical analysis using GraphPad Prism software version 8.0.0. Statistical significance was assessed using Student’s T-test. The significance level of *p* < 0.05 was used to determine statistical significance across all analyses.

### 2.4. Photosynthetic pigments

Chlorophyll *a* and chlorophyll *b* were extracted at 31 DAI using dimethyl sulfoxide (DMSO). Two rosette discs from the fifth leaf were incubated in 1 mL of DMSO for 72 h at room temperature in the dark. Absorbance was measured at 480, 649.1, and 665.1 nm (NanoDrop™ 2000), and pigment concentrations were calculated based on standard equations (Ca = 12.74 × A665.1 − 3.62 × A649.1; Cb = 25.06 × A649.1 − 6.5 × A665.1). Results were expressed as μg/g fresh weight. Statistical analysis was based on the Student’s T-test for all comparisons. Statistical significance was defined as *p*-value < 0.05.

### 2.5. RNA extraction and Real-time quantitative PCR

For molecular analyses, plant roots were macerated in liquid nitrogen and immediately stored at −80°C. At 11 DAG, total RNA was extracted from inoculated and non-inoculated samples of each genotype using a lithium chloride–based protocol, as described by (Logemann *et al*., 1987). RNA concentration was determined with a NanoDrop™ 2000c spectrophotometer (Thermo Scientific, USA), and its integrity was assessed by electrophoresis in a 1% agarose gel stained with ethidium bromide. Approximately 5 µg of total RNA was treated with DNase I (Biolabs), according to the manufacturer’s instructions, and used for the synthesis of the first strand of cDNA with GoScript reverse transcriptase (Promega). Gene expression analyses were performed by RT-qPCR using SYBR Green PCR Master Mix (Applied Biosystems). Each reaction was assembled in a final volume of 10 µL, containing 2 µL of cDNA diluted 1:3, 5 µL of SYBR Green mix, 1 µL of each primer (forward and reverse, 10 µM) and 1 µL of sterile ultrapure water. The primers specific for the target genes analyzed are listed in Table S1. Expression normalization was performed based on the constitutive genes *UBIQUITIN 14* (*UBI14*) and *GLYCERALDEHYDE-3-PHOSPHATE DEHYDROGENASE* (*GAPDH*) of *A. thaliana*. Data analysis followed the ΔCT method, based on the subtraction of the CT value of the constitutive gene in relation to the target gene. All experiments were conducted with three biological replicates per condition.

### 2.6. DNA extraction and Real time quantitative PCR absolute bacterial quantification

Total DNA extraction was performed using CTAB 2% protocol (Doyle and Doyle, 1987). DNA concentration was determined with a NanoDrop™ 2000c spectrophotometer, and its integrity was assessed by electrophoresis in a 1% agarose gel stained with ethidium bromide. Total DNA was treated with 0,01g/mL RNase A and incubated at 37 °C for 1 h. To determine the bacterial cell number cells (cells g⁻¹ plant tissue), the average Ct value from two technical replicates for each DNA sample was used to quantify *G. diazotrophicus* PAL5, *H. seropedicae* HRC54, *A. brasilense* Ab-V6. Genus-specific standard curve equations, previously established in the laboratory, were applied for quantification. The number of cells were inferred from the amplicon copy number, assuming the target sequence is single copy per genome. Target-specific primer pairs are listed in Table S1. For each treatment, cells per gram of tissue were calculated from the DNA samples using the following equation: Absolute number of cells = A × B (ng μL⁻¹) × C (μL) / D (ng) × E (g). Where, A = number of fragment copies, obtained from the strain-specific standard curve; B = Concentration of extracted genomic DNA; C = volume of DNA extracted; D = amount of DNA used in the qPCR reaction; E = initial plant tissue mass used for DNA extraction.

### 2.7. Metagenomic Barcode Analysis

Plants used in the metagenomic barcode analysis experiment were grown under the low nutrient condition protocol and collected at 20 DAG. For each genotype Col-0 wild type and *aip10-1* plants, one sample pool consisting of three plants was collected. Metagenomic barcode analysis experiment was performed exclusively from the root tissue of each genotype. Soil samples were obtained as a sample pool composed of material from the three pots corresponding to the plants whose roots were collected.

Total DNA was extracted separately from root and rhizosphere soil samples of *A. thaliana* Col-0 and the mutant *aip10-1* using a commercial soil DNA extraction kit. The V3–V4 regions of the bacterial 16S rRNA gene were amplified with universal primers containing Illumina adapters, and amplicon libraries were prepared following standard protocols. Sequencing was performed on an Illumina MiSeq platform (PE 500 bp). Taxonomic assignment was performed against the SILVA 138 database, with removal of non-bacterial sequences. Relative abundance of bacterial taxa was calculated separately for root and soil samples, and results were visualized using GraphPad Prism 10.5.0 for bar plots and comparisons between genotypes. Heatmaps of selected PGPB, diazotrophs, and pathogens were generated in R (pheatmap package), based on log-transformed relative abundances and hierarchical clustering.

### 2.8. Confocal microscope and microscopic analysis

Roots of *A. thalian*a ecotype Col-0 wild type and *aip10-1* inoculated with *G. diazotrophicus* Pal5-RFP and *H. seropedicae* RAM10-GFP were analyzed 7 DAI. Imaging was conducted using a Leica TCS SPE confocal microscope. Plant samples were mounted on microscope slides with a drop of water and covered with a coverslip for observation under a light microscope. For RFP observation (from *G. diazotrophicus* Pal5-RFP), samples were excited at 568 nm and emission was collected at 640 nm. For *H. seropedicae* RAM10-GFP (BR 11335), samples were excited at 475 nm and emission was collected at 509 nm. Differential interference contrast (DIC) images were also captured simultaneously using bright field. Image acquisition and processing were performed using Leica LAS AF software.

## 3. Results

### AIP10 expression profile in response to plant growth-promoting bacteria

To explore the relationship between PGPB association, plant genotype, and *AIP10* expression, we examined the transcriptional profile of the sugarcane (*Saccharum spp.*) *AIP10* homologue based on previously published RNA-seq datasets from two genotypes differing in their capacity to associate with beneficial bacteria and to sustain BNF: SP70-1143 (high BNF) and Chunee (low BNF) (Urquiaga and Boddey, 1992; Carvalho *et al*., 2022). Transcript levels were analyzed in root and shoot tissues from plants grown under natural bacterial colonization and bacteria-free hydroponic conditions. Fold change values (SP70-1143/Chunee) were calculated based on RPKM values to compare the genotype with higher association efficiency with diazotrophic bacteria to the less efficient genotype.

Under natural colonization conditions, the *AIP10* homolog exhibited lower expression levels in both roots and shoots of the high-BNF genotype SP70-1143 relative to Chunee (Fig. 1A). In contrast, under bacteria-free conditions (BF), *AIP10* expression was upregulated in both tissues root and shoot of SP70-1143. These results indicate that *AIP10* expression is reduced in the genotype exhibiting higher BNF capacity when associated with beneficial bacteria, supporting its role as a negative regulator of plant growth, and suggesting its modulation may participate in growth promotion in response to beneficial plant-bacteria interaction.

**Figure 1.**
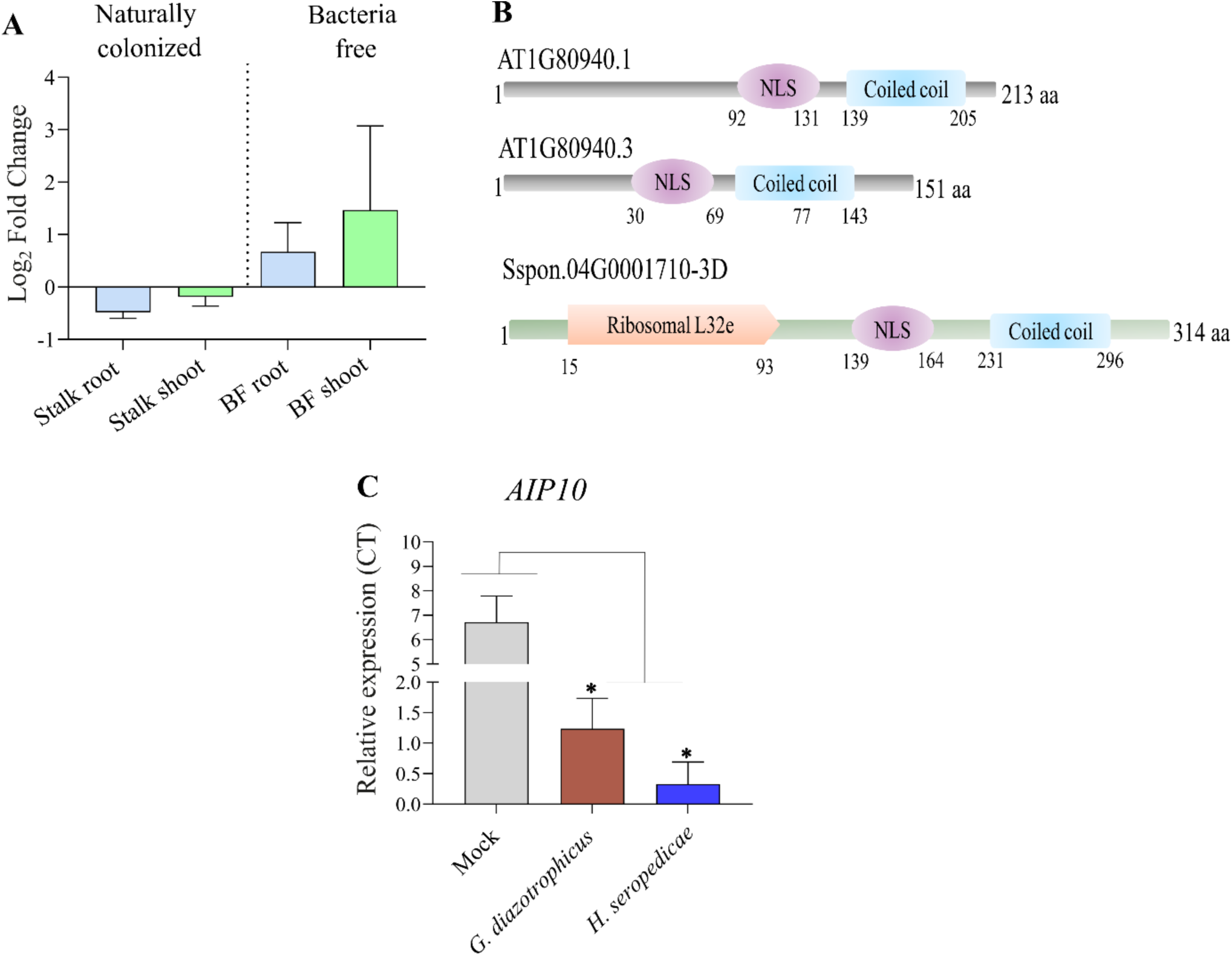
Conserved modulation of *AIP10* expression by plant growth–promoting bacteria and its nuclear protein structure in *A. thaliana* and sugarcane. (A) Expression profile of the *AIP10* homolog in sugarcane root and shoot tissues of two BNF contrasting genotypes (SP70-1143, high BNF; Chunee, low BNF) from plants naturally colonized by PGPB (Stalk) and plants grown under bacteria-free conditions (BF). Expression values are shown as log_2_ fold change. **(B)** Predicted protein structures of *AIP10* putative homologs encoded by two isoforms *AT1G80940.1* and *AT1G80940.3* from *A. thaliana*, and by *Sspoon.04G0001710-3D* from *Saccharum spontaneum*. Protein domains were predicted using the SMART program. The scheme shows the nuclear localization signal (NLS), coiled-coil, and ribosomal L32e domains, suggesting a possible role related to nuclear regulation and protein interaction, potentially conserved between species. **(C)** Expression levels of *AIP10* in *A. thaliana Col-0* plants at 7 DAI after inoculation with two different PGPB strains, *G. diazotrophicus* and *H. seropedicae*.

A comparative protein domain analysis revealed that AIP10 proteins from *A. thaliana* and sugarcane share a conserved domain architecture, including predicted nuclear localization signals (NLS) and coiled-coil domains (Fig. 1B). This high level of structural conservation suggests a conserved regulatory function of AIP10 across monocot and dicot species. In agreement with these findings, the data revealed a similar *AIP10* expression profile in both plant species, suggesting that these proteins may share similar biological functions. Comparative protein domain analysis of two AIP10 orthologous isoforms in *A. thaliana* (AT1G80940.1 and AT1G80940.3) and one putative isoform in *S. spontaneum* (Sspoon.04G0001710-3D) further confirmed the conservation of nuclear localization and coiled-coil domains, while revealing the presence of an additional ribosomal L32e domain in the sugarcane protein.

Next, we investigated the modulation of *AIP10* expression profile by inoculation with beneficial PGPB in Col-0 plants from *A. thaliana*. Plant material was collected at 7 DAI with the diazotrophic PGPB *H*. *seropedicae* or *G. diazotrophicus,* to assess the *AIP10* transcriptional response during the early stages of plant–microbe interaction (Fig. 1C). The results showed a significant repression of *AIP10* expression in response to both bacterial strains compared to mock-treated controls, with a stronger down-regulation observed upon inoculation with *H. seropedicae* (Fig. 1C). These data suggest that *AIP10* expression is reduced during early stages of plant-PGPB interactions. Together, the results indicate that AIP10 displays a conserved domain architecture across species, while its expression is differentially modulated by bacterial colonization in sugarcane genotypes contrasting for BNF efficiency and in *A. thaliana*.

To explore molecular pathways involved in plant–microorganism interactions potentially modulated by *AIP10* repression, we reanalyzed previously generated RNA-seq data in the root transcriptomes of *A. thaliana* Col-0 and *aip10-1* at 11 DAG (Montessoro *et al*., 2025). We examined enriched functional categories across metabolic pathways related to Cell wall formation, Response to stimuli, Immune response to bacteria, Responses to hormones, Responses to lipids, Carbohydrates and General metabolic Processes (Fig. S1).

In cell wall–related categories, genes directly involved in cell wall reinforcement and stress-associated remodeling were downregulated (Fig. S1A), including *EXT15* (*Extensin 15*, AT1G23720), *LRX3* (*Leucine-rich repeat extensin 3*, AT4G13340) and LRX4 (At3g24480. The repression of extensins and LRX proteins, which are essential for cell wall assembly, integrity, and mechanical strength, suggests increased cell wall plasticity that may facilitate bacterial association and root tissue colonization (Draeger *et al*., 2015; Kandel *et al*., 2017). A distinct group of proteins associated with cell wall expansion and extracellular matrix organization was significantly induced at 11 DAG, including *GASA1* (Gibberellic Acid Stimulated Arabidopsis 1, AT1G75750) and *LCR69* (Low-molecular-weight Cysteine-rich protein 69, AT2G02100), suggesting enhanced wall loosening and elongation processes. Consistent with this pattern of cell wall modification, genes encoding enzymes involved in polysaccharide modification and pectin remodeling were also upregulated, including *PME46* (Pectin Methylesterase 46, AT5G04960), *XHT* (Xyloglucan endotransglucosylase/hydrolase, AT4G25820), and *ATTIP2* (Tonoplast Intrinsic Protein 2, AT3G16240). These changes suggest active restructuring of the polysaccharide network that regulates wall extensibility and porosity in *aip10-1* plants, potentially enhancing apoplastic permeability and promoting cell-to-cell communication (Dora *et al*., 2022).

In the hormone-responsive category, the auxin-responsive genes *SAUR69* (Small Auxin Upregulated RNA 69) and *SAUR72* (Small Auxin Upregulated RNA 72) were strongly upregulated, suggesting a role in promotion of plant growth. Up-regulation of *RSL4* (*bHLH85*, ROOT HAIR DEFECTIVE 6-LIKE 4), a transcription factor, that positively regulates root hair development, was upregulated in *aip10-1* roots. Enhanced root hair growth is a commonly reported response to inoculation with PGPB (Zhao et al., 2025).

Some genes involved in defense or immunity pathways associated with bacterial interactions were repressed, whereas others were upregulated in *aip10-1* roots, consistent with the regulatory adjustments required for beneficial associations with PGPB (Carvalho et al, 2016; Stringlis *et al*., 2018). Among the most downregulated genes were *CYP71A12* and *CYP71A13* (Cytochrome P450 71A12 and– Cytochrome P450 71A13), which are key enzymes involved in camalexin and tryptophan-derived secondary metabolite biosynthesis. Their repression suggests a reduction in defense pathways. In contrast, a distinct subset of bacterium-responsive genes was strongly induced, reflecting activation of regulatory pathways associated with plant–microbe communication and systemic signaling. A subset of stimulus-related genes, including *PDF2.2* (*Plant Defensin 2.2*), *ATL15* (*RING-type E3 ubiquitin ligase 15*), AT1G22770, and AT1G15010, was induced, suggesting the activation of regulatory and antimicrobial components that may contribute to controlling the number of bacteria that could associate with and colonize the root tissue. The most strongly induced gene was *ILA* (ILITHYIA), which is required for plant immunity and is directly associated with basal defense in *A. thaliana* against the pathogenic bacteria *Pseudomonas syringae* (Monaghan and Li, 2010). It also plays a central role in coordinating developmental and immune signaling (Faus *et al*., 2018).

Regulation of metabolic, lipid, and carbohydrate-responsive genes in *aip10-1* revealed coordinated transcriptional reprogramming affecting primary metabolism and stress-associated pathways. Repression of *PAD3* (Phytoalexin Deficient 3) and *CBP60G* (Calmodulin-Binding Protein 60G) supports an attenuation of defense-related metabolic pathways, indicating a general downregulation of stress-associated programs. Within this metabolic shift, genes involved in carbohydrate-related processes also showed regulatory adjustments. In particular, repression of *GLS1* (Glucan Synthase-Like 1) together with induction of *ATL15* (RING-Type E3 Ubiquitin Ligase 15) suggests a reprogramming of carbon-associated regulation, reflecting an adjustment of carbohydrate metabolism rather than a simple reduction in carbon availability.

Together, the transcriptional changes observed in *AIP10*-silenced plants revealed modulation of core metabolic processes coupled with regulation of defense and growth-related pathways. These changes are consistent with transcriptional reprogramming associated with plant responses to beneficial endophytic or associative PGPB and aligned with the effects of reduced *AIP10* expression observed under natural bacterial colonization in sugarcane.

### Root-associated microbiota composition in A. thaliana wild-type and aip10-1 plants

Given that transcriptional reprogramming in *aip10-1* plants affected multiple pathways related to plant–microbe interactions, we next examined whether AIP10 silencing altered the composition of the root-associated microbiota in *A. thaliana*. A metagenomic barcode analysis of the root-associated bacterial communities and the rhizosphere soil of non-inoculated Col-0 and *aip10-1* plants was performed.

The Venn diagram revealed a complex distribution of shared and unique taxa among the four sample types, comprising soil and roots from both genotypes. Col-0 and *aip10-1* roots each harbored a high number of genotype-specific bacterial species (237 and 203, respectively) (Fig. 2A), indicating distinct recruitment patterns. Despite these differences, a substantial core microbiome of 975 bacterial species was shared across all samples, highlighting a conserved microbial baseline between soil and plant genotypes. Additionally, soil samples associated with Col-0 and *aip10-1* each contained unique taxa (173 and 220 species, respectively) (Fig. 2A), reflecting environmental and genotype-driven filtering effects. Together, these findings demonstrate that *AIP10* silencing reshapes root-associated bacterial communities while maintaining a large portion of shared microbial taxa.

**Figure 2.**
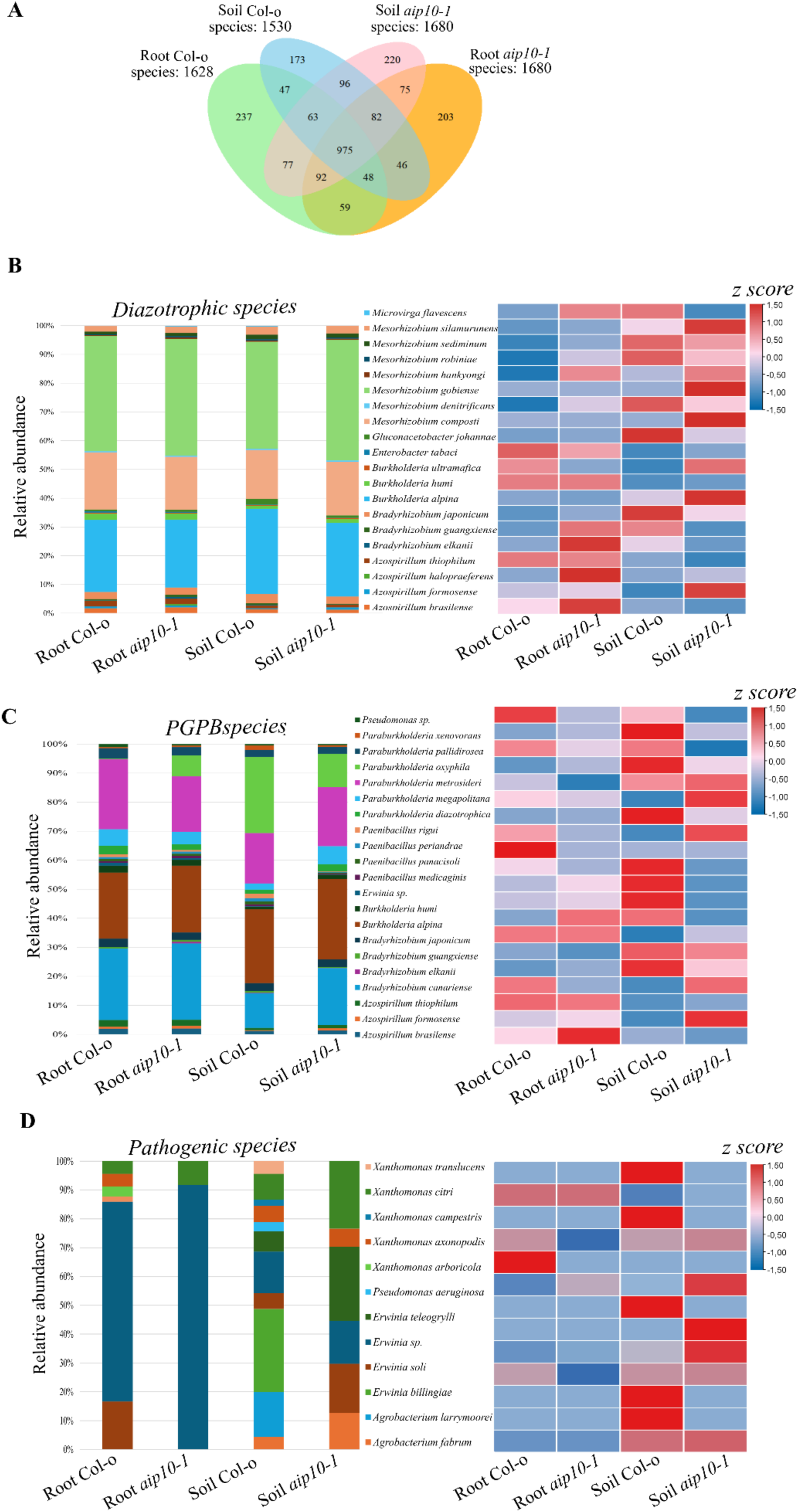
Taxonomic distribution and enrichment patterns of key microbial functional groups in roots and bulk soil of non-inoculated Col-0 and *aip10-1* plants. (A) Venn diagram shows shared and unique microbial species associated with roots and surrounding soil from Col-0 and *aip10-1* plants. **(B–D)** Relative abundance (left panels) and z-score heatmaps (right panels) for bacterial groups with functional relevance to plant performance, including **(B)** potential diazotrophic bacterial species, **(C)** potential plant growth-promoting bacteria (PGPB), and **(D)** potential pathogenic bacterial species. Data are shown for root and soil compartments of Col-0 wild type and *aip10-1* plants. In the heat map, red represents higher abundance, and blue represents lower abundance, according to the z-score value.

For a more focused assessment of functionally relevant components of the native microbiome potentially associated with growth promotion and innate immune responses in the host plant, the abundance of selected bacterial categories was analyzed by z-score values (Table S2), including potential diazotrophic bacteria, PGPB, and plant pathogenic bacteria.

Comparative profiling of bacterial communities associated with the plant root compartment revealed pronounced differences between Col-0 and *aip10-1* genotypes (Fig. 2B-C). At the species level, roots of *aip10-1* plants exhibited increased relative abundance of several potential diazotrophic bacteria, including, *Azospirillum brasilense* (z-score 1,4), *Azospirillum halopraferens* (z-score 1,5), *Bradyrhizobium elkanii* (z-score 1,4) and *Bradyrhizobium guangxiense* (z-score 0,9). In addition, plant PGPB species were enriched in *aip10-1* roots in comparison with Col-0 roots, as *Paenibacillus medicaginis* (z-score 0,9) (Fig. 2C).

In contrast, roots of Col-0 plants displayed higher relative abundance of several pathogenic species, compared to *aip10-1* roots (Fig. 2D). Taxa such as *Xanthomonas arboricola* (z-score 1,5), *Xanthomonas axonopodis* (z-score 0,5), *Pantoea brenneri* (z-score 1,5), and *Erwinia soli* (z-score 0,4) were comparatively more abundant in Col-0 and were not detected in *aip10-1*. These data suggest that *aip10*-1 plants tend to preferentially recruit and associate with specific beneficial PGPB species in the root compartment while being depleted in potentially pathogenic taxa.

A similar pattern was observed in soil-associated bacterial communities (Fig. 2B-C). Soils cultivated with *aip10-1* plants were characterized by increased relative abundance of several potential diazotrophic species, including *Mesorhizobium composti* (z-score 31,2), *Mesorhizobium gobiense* (z-score 52,7), *Mesorhizobium hankyongi* (z-score 1,0), *Mesorhizobium silamurunense* (z-score 3,7) *Azospirillum formosense* (z-score 1,5), *Burkholderia alpina* (z-score 6,3), *Burkholderia ultramafica* (z-score 0,7), when compared with soils from Col-0 plants. Likewise, PGPB species such as *Paraburkholderia diazotrophica* (z-score 1,1), *Paraburkholderia megapolitana* (z-score 1,2), *Paraburkholderia metrosideri* (z-score 0,9) and *Bradyrhizobium canariense* (z-score 0,9) were more abundant in soils associated with *aip10-1*.

On the other hand, soils from Col-0 plants harbored higher relative abundances of potential pathogenic species (Fig. 2D), including *Xanthomonas campestris* (z-score 1,5), *Xanthomonas translucens* (z-score 1,5), *Erwinia billingiae* (z-score 1,5), and *Agrobacterium larrymoorei* (z-score 1,5). Together, these results suggest that the loss of AIP10 function reshapes the root-associated bacteria in roots and rhizospheric soil, increasing the prevalence of beneficial bacteria while reducing potential pathogens.

### Efficiency of PGPB bioinoculant colonization in aip10-1 plants

As *AIP10* silenced mutants showed increased abundance of beneficial bacteria in their root microbiome, we next investigated whether *aip10-1* mutants exhibited enhanced colonization in response to inoculation with diazotrophic PGPB bioinoculants, compared to wild-type Col-0 plants, by quantifying and assessing the dynamics of bacterial root colonization.

Initially, an *in vitro* inoculation system followed by greenhouse cultivation was established to study the interaction of *aip10-1* mutant plants with two N_2_-fixing bacterial strains, *H. seropedicae* HRC54 or *G. diazotrophicus* Pal5. Representative images of *aip10-1* plants at 20 DAG, inoculated with either strain, are shown in Fig. 3A. Under *in vitro* growth conditions, root architectural parameters were evaluated revealing that inoculated *aip10-1* plants displayed a significant increase in primary root length compared with mock-treated plants and inoculated Col-0 controls (Fig. 3B). In addition, the number of lateral roots was significantly higher in inoculated *aip10-1* plants, particularly in response to PAL5 (Fig. 3C).

**Figure 3.**
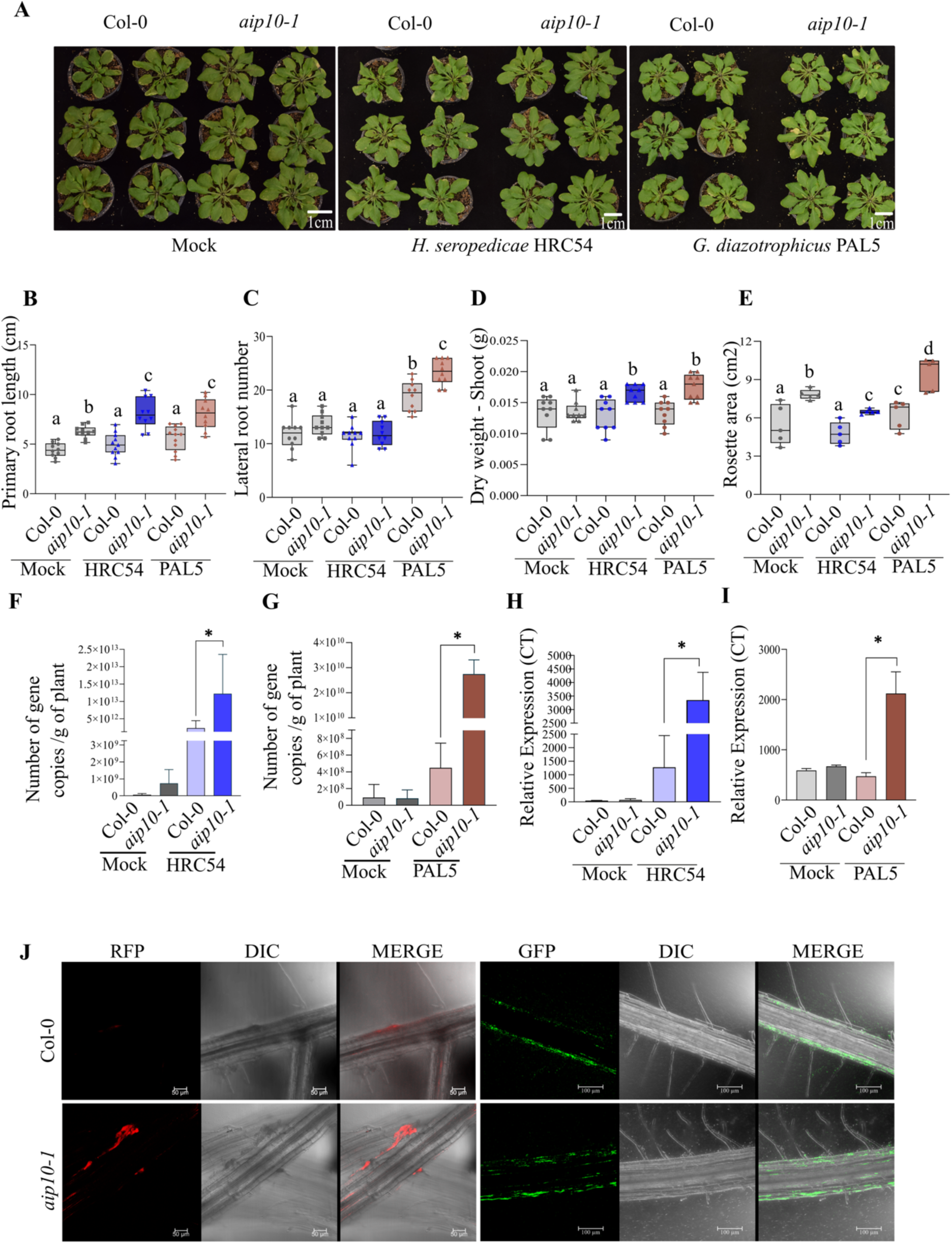
Functional analyses of *AIP10* silencing in *A. thaliana* root colonization and association with diazotrophic PGPB strains *H. seropedicae* HRC54 and *G. diazotrophicus* Pal5. (A) Representative images of Col-0 and *aip10-1* plants mock-treated, inoculated with *H. seropedicae* HRC54 or *G. diazotrophicus* Pal5 at 20 days after germination (DAG). Scale bars: 1cm. *Note that images in the three panels have different Bars.* (B–E) Growth and developmental parameters of Col-0 and *aip10-1* plants inoculated with *H. seropedicae* HRC54 or *G. diazotrophicus* Pal5. **(B)** Primary root length and **(C)** lateral root number measured at 15 DAG, the box plot graphs show the distribution of data of 10 individuals. **(D)** Dry weight of the shoots, using nine biological replicates, and **(E)** rosette area, using five biological replicates, measured at 21 DAG. Data are presented as box plots showing median, interquartile range, and minimum/maximum values. **(F, G)** Absolute qPCR quantification of *H. seropedicae* HRC54 and *G. diazotrophicus* Pal5 colonization in roots of Col-0 and *aip10-1* at 7 DAI, respectively. The data represents the number of bacterial cells per gram of root tissue, obtained by absolute qPCR quantification. Bars represent mean ± standard deviation of three biological replicates and each biological replicate (pool of five plants) was analyzed with three technical replicates. **(H, I)** Relative expression of the *nifH* gene in *H. seropedicae* HRC54 and *G. diazotrophicus* Pal5 colonizing Col-0 and *aip10-1* roots at 7 DAI, quantified by RT-qPCR. Gene expression values are shown as fold-change relative to GAPDH and UBI14 reference genes. Bars represent mean ± standard deviation of the relative mRNA expression in three biological replicates and each biological replicate (pool of five plants) was analyzed with three technical replicates. **(J)** Representative confocal microscopy images showing *G. diazotrophicus* (RFP-labeled) and *H. seropedicae* (GFP-labeled) colonization patterns in Col-0 and *aip10-1* roots at 11 DAG and 7 DAI. Images include RFP, GFP, DIC, and merged channels. Scale bars: 50 μm. The box plot graphs represent the interquartile range (IQR), with the inner line running down the median. The whiskers extend to the maximum and minimum values, including all points within that range. The differences between groups in all analyses were confirmed by One-Way ANOVA (p < 0.05), and different letters indicate statistically different means according to Tukey’s test at 5% probability. DAG, Days After Germination. Statistical analyses were performed using Student’s t-test at a 5% significance level. (*) indicates statistically significant differences (p ≤ 0.05).

To determine whether these early root architectural changes translate into later developmental outcomes, shoot-related traits were assessed at later developmental stages. At 21 DAG, *aip10-1* plants inoculated with both bacterial strains accumulated significantly higher shoot biomass (Fig. 3D) and developed larger rosette areas compared with Col-0 plants and mock-treated controls (Fig. 3E).

Quantification of bacterial colonization based on absolute values of single-copy bacterial gene using genus-specific primers showed that *aip10-1* roots of inoculated plants were better colonized by both bioinoculants, compared to wild-type Col-0 roots (Fig. 3F and G). To evaluate if the higher colonization by *G. diazotrophicus* or *H. seropedicae* was correlated with actively metabolic bacteria, *nifH* mRNA expression was analyzed. The *nifH* is present in most diazotrophic microorganisms and encodes a key component of nitrogenase, making it an ideal marker for assessing the stimulation of N_2_-fixation metabolism in bacterial communities (Nichio *et al*., 2025). As shown in Fig. 3H and I, PGPB-inoculated *aip10-1* plants showed significantly higher relative expression of *nifH* than inoculated wild-type plants during association with both HRC54 or PAL5, respectively.

To investigate whether *aip10-1* mutants affect the niche of bacterial colonization in root tissues, RFP- and GFP-labeled G. *diazotrophicus* and *H. seropedicae* bacteria, respectively, were used as bioinoculants and monitored by confocal microscopy. The data showed differences in Col-0 and *aip10-1* roots in relation to the pattern of bacterial colonization. In mock Col-0 roots, both RFP- and GFP-labeled bacteria showed minimal and discontinuous fluorescence, indicating low colonization levels. In contrast, inoculated *aip10-1* roots displayed strong and spatially extended fluorescent signals, with well-defined aggregates and continuous bands along the root surface (Fig. 3J). A detailed analysis of distinct root regions further demonstrated that these differences were spatially structured. In the meristematic zone, both genotypes exhibited low colonization (Fig. S2). However, beginning in the root hair zone, *aip10-1* showed a marked increase in bacterial attachment, with continuous fluorescent bands extending along the root axis. This pattern became more evident in the elongation zone, where large portions of the cortex and epidermis were covered by bacteria, in sharp contrast to the fragmented and sparse pattern observed in Col-0 (Fig. S2). The highest colonization levels were observed at lateral root emergence sites, where inoculated aip*10-1* roots displayed extensive and well-defined bacterial aggregates, whereas Col-0 presented only small, scattered foci. Together, these spatial colonization patterns indicate that loss of AIP10 enhances root colonization, particularly in regions of active growth and lateral root formation, resulting in greater density, continuity, and stability of bacterial association.

Together, these results indicate that *AIP10* silencing is associated with enhanced bacterial colonization and activity in roots, accompanied by pronounced changes in root architecture and shoot growth following inoculation.

### Molecular mechanisms of plant growth promotion in PGPB inoculated aip10-1 plants

As *AIP10* silenced mutants showed increased root colonization by beneficial bacteria and enhanced plant growth promotion, compared to wild-type Col-0 plants, we investigated the molecular mechanisms that may contribute to the improved responsiveness of *aip10-1* plants to PGPB bioinoculants.

First, we investigated the modulation of cell division controls in inoculated *aip10-1* and Col-0 plants. AIP10 has previously been described as a negative modulator of cell division, through its interaction with ABAP1 and regulation of ABAP1 target gene expression (Montessoro *et al*., 2025). Also, *ABAP1* expression is negatively regulated in *aip10-1* mutants. Therefore, the expression levels of *ABAP1* and *CDT1a*, a target gene negatively regulated by ABAP1 and a positive marker of DNA replication, were evaluated by RT-qPCR (Fig. 4). In *aip10-1* plants, *ABAP1* expression was more strongly repressed following inoculation with PAL5 and HRC54 compared to mock-treated mutants, whereas inoculated Col-0 plants consistently exhibited higher *ABAP1* expression than *aip10-1* mutants (Fig. 4A). Accordingly, *CDT1a* expression was induced in *aip10-1* plants in response to both beneficial inoculations, with PAL5 triggering significantly higher induction than in mock-treated plants (Fig. 4B). Expression of the cell division marker *CYCB1;1* was strongly induced in inoculated *aip10-1* mutant plants compared to mock-treated controls, suggesting that the increased DNA replication rates stimulated by beneficial bacterial colonization were followed by higher cell division rates in *aip10-1* plants (Fig. 4C). Overall, increased colonization by PGPB bioinoculants in *aip10-1* mutant plants enhanced the transcriptional regulation of ABAP1 and associated pathway genes, consistent with the greater growth promotion observed following inoculation.

**Figure 4.**
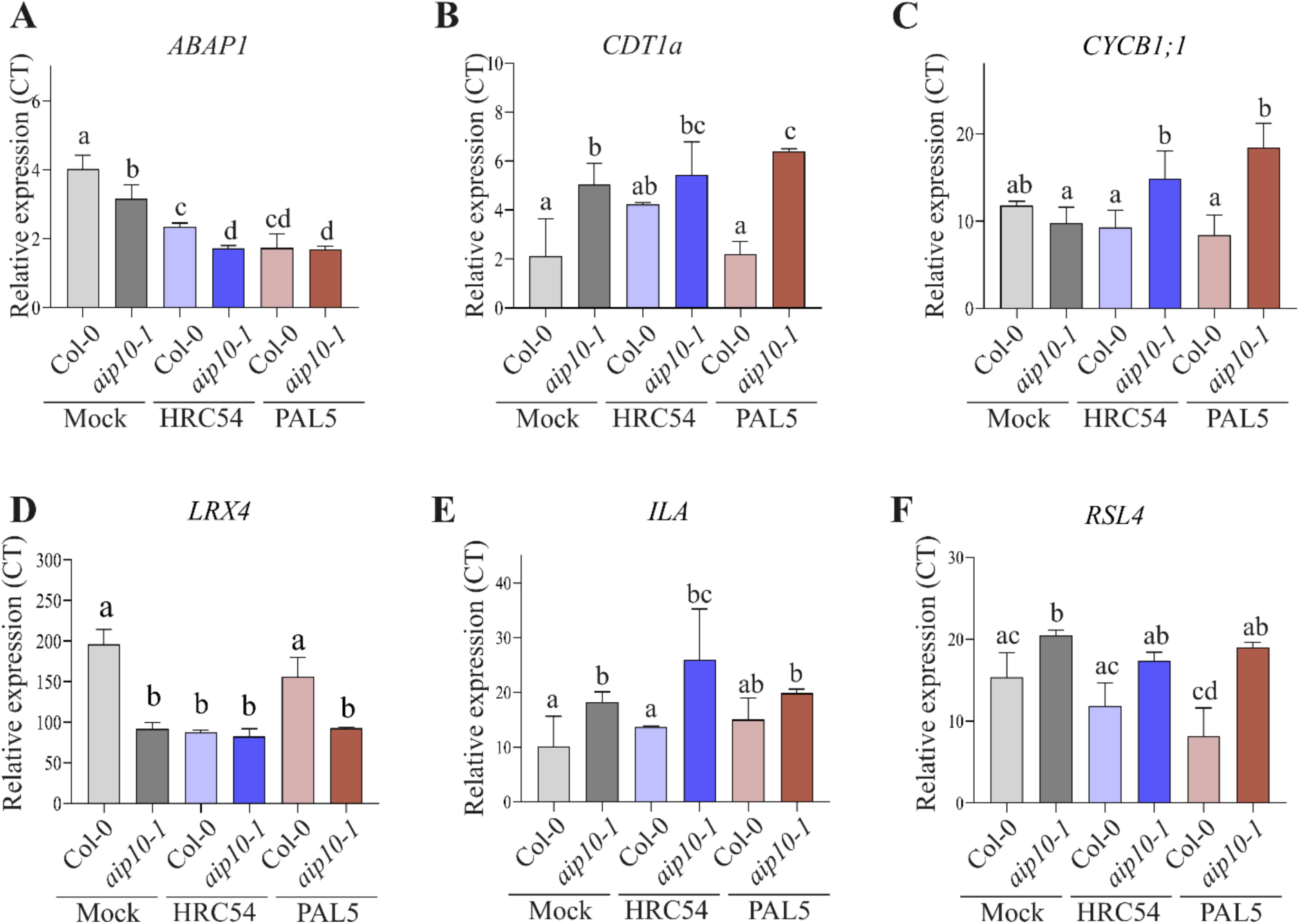
RT-qPCR analysis of genes involved in cell division (ABAP1 regulatory network), cell wall organization, bacterial response, and root morphogenesis in *aip10-1* and wild-type Col-0 roots, at 7 DAI. (A) *ABAP1*; (B) *CDT1a*; (C) *CYCB1;1*; (D) *LRX4*; (E) *ILA* and (F) *RSL4*. RT-qPCR results are presented as the ratio of expression of each gene relative to the constitutive expression of *GAPDH* and *UBI14* reference genes when compared to mock and inoculated plants. Bars represent mean ± standard deviation of the relative mRNA expression in three biological replicates and each biological replicate analyzed with three technical replicates. Statistical analysis was performed using the Student’s t-test (P ≤ 0.05). Means with different letters are significantly different at 5% level of confidence.

Next, based on the RNAseq data from non-inoculated Col-0 and *aip10-1* plants, three genes related to cell wall organization, defense against bacteria, or root hair development were selected for expression analysis in roots under beneficial inoculation, by RT-qPCR (Fig. S1). According to transcriptomic data, LRX*4* was downregulated in *aip10-1* compared to Col-0. This expression pattern was confirmed in mock-treated *aip10-1* plants; however, no additional changes were observed upon bacterial inoculation when compared to wild-type plants (Fig. 4D). Similarly, the upregulated expression of *ILA* in *aip10-1* roots identified in the transcriptome analysis was confirmed by RT-qPCR, and similar *ILA* levels were observed in inoculated plants, compared to Col-0 (Fig. 4E). Transcriptomic data revealed upregulation of *RSL4* in *aip10-1* roots. This result was confirmed by RT–qPCR, which showed higher expression levels in mock-treated *aip10-1* plants compared to Col-0, with expression remaining elevated in the presence of both bacterial strains (Fig. 4F). Our analyses showed that gene expression profiles were largely similar between non-inoculated and inoculated *aip10-1* plants, including genes related to cell wall dynamics and bacterial responses.

The absence of further transcriptional modulation upon PGPB inoculation suggested that *aip10-1* mutants may already be transcriptionally primed for enhanced association with beneficial plant–microorganism interactions. This genotype appears to provide physiological conditions that favor colonization by beneficial and diazotrophic bacteria, thereby increasing its capacity to respond to their growth-promoting effects.

### Response of *aip10-1* plants to diazotrophic PGPB inoculation under high and low fertilization conditions

Given the enhanced association between *aip10-1* plants and PGPB bioinoculants, together with increased *nifH e*xpression indicating higher nitrogenase activity in inoculated *aip10-1* roots, we investigated whether PGPB inoculation could reduce the dependence of *aip10-1* mutant plants on mineral fertilization. To address this question, inoculation responses of *aip10-1* and Col-0 plants were analyzed under conditions of low and high nutrient availability.

The *in vitro* inoculation system was used, followed by greenhouse cultivation under low (0.06 g/L NPK 20:20:20, corresponding to 12 mg/L N, 5.2 mg/L P and 10 mg/L K) and high (0.12 g/L NPK 20:20:20, corresponding to 24 mg/L N, 10.4 mg/L P and 20 mg/L K) nutrient availability. Col-0 and *aip10-1* mutant plants were inoculated with the PGPB *H. seropedicae* HRC54, or *G. diazotrophicus* Pal5, or a commercial bioinoculant (Biofree) containing the strains *A. brasilense* Ab-V6 and *P. fluorescens* CCTB03. Four treatment groups were established for both Col-0 and *aip10-1* plants: i) mock-treated plants grown under low fertilization, ii) mock-treated plants grown under high fertilization, iii) plants inoculated with HRC54 or PAL5 or Biofree grown under low fertilization, iv) plants inoculated with HRC54 or PAL5 or Biofree grown under high fertilization (Fig. S3). First, root colonization with the PGPB bioinoculants (HRC54 or Pal5 or Biofree) was validated in plants at 7 DAI, before starting differential fertilization treatments, through absolute quantification of a single-copy bacterial gene using genus-specific primers by qPCR (Fig. 3F-G and Fig. S4). Bacterial counts showed that *aip10-1* roots were colonized by a higher number of bacterial cells per gram of plant tissue compared to Col-0, suggesting enhanced interaction between mutant plants and the bioinoculants.

The effects of fertilization with high or low doses of nutrients were analyzed in the shoots and roots at 35 DAG (31 DAI) (Fig. 5A and B). Our results showed that *aip10-1* plants were more responsive to bioinoculants under low nutrient availability, exhibiting enhanced growth, like the phenotype observed in mock-treated mutant plants grown under high nutrient availability. In shoot, this beneficial effect was observed through analyses of rosette area and dry biomass, where *aip10-1* inoculated plants presented 22% (HRC54) (Fig. 5C) and 26% (Pal5) (Fig. 5F) larger area, as well as 33% (HRC54) (Fig. 5D) and 35% (Pal5) (Fig. 5G) greater dry weight when grown under low N availability compared to Col-0 plants under the same conditions. Furthermore, rosette area and dry biomass increased by 30% (Fig. 5C and F) and 42% (Fig. 5D and G), respectively, in mock-treated *aip10-1* mutant plants under conditions of high nutrient availability. Chlorophyll content analyses demonstrated this same pattern, with an increase in response to both inoculations in *aip10-1* plants compared to wild-type plants under low fertilization conditions (Fig. 5E and H), consistent with the increase observed in mock mutant plants grown under high fertilization, corroborating previous results. Parallel experiments were conducted using a commercial bioinoculant, Biofree (Fig. S4). Under low nutrient availability, *aip10-1* plants inoculated with Biofree exhibited a marked increase in rosette expansion and biomass accumulation compared to mock-treated mutants (Fig. S4). Again, this growth promotion was comparable to that observed in mock-treated *aip10-1* plants grown under high fertilization conditions (Fig. S4).

**Figure 5.**
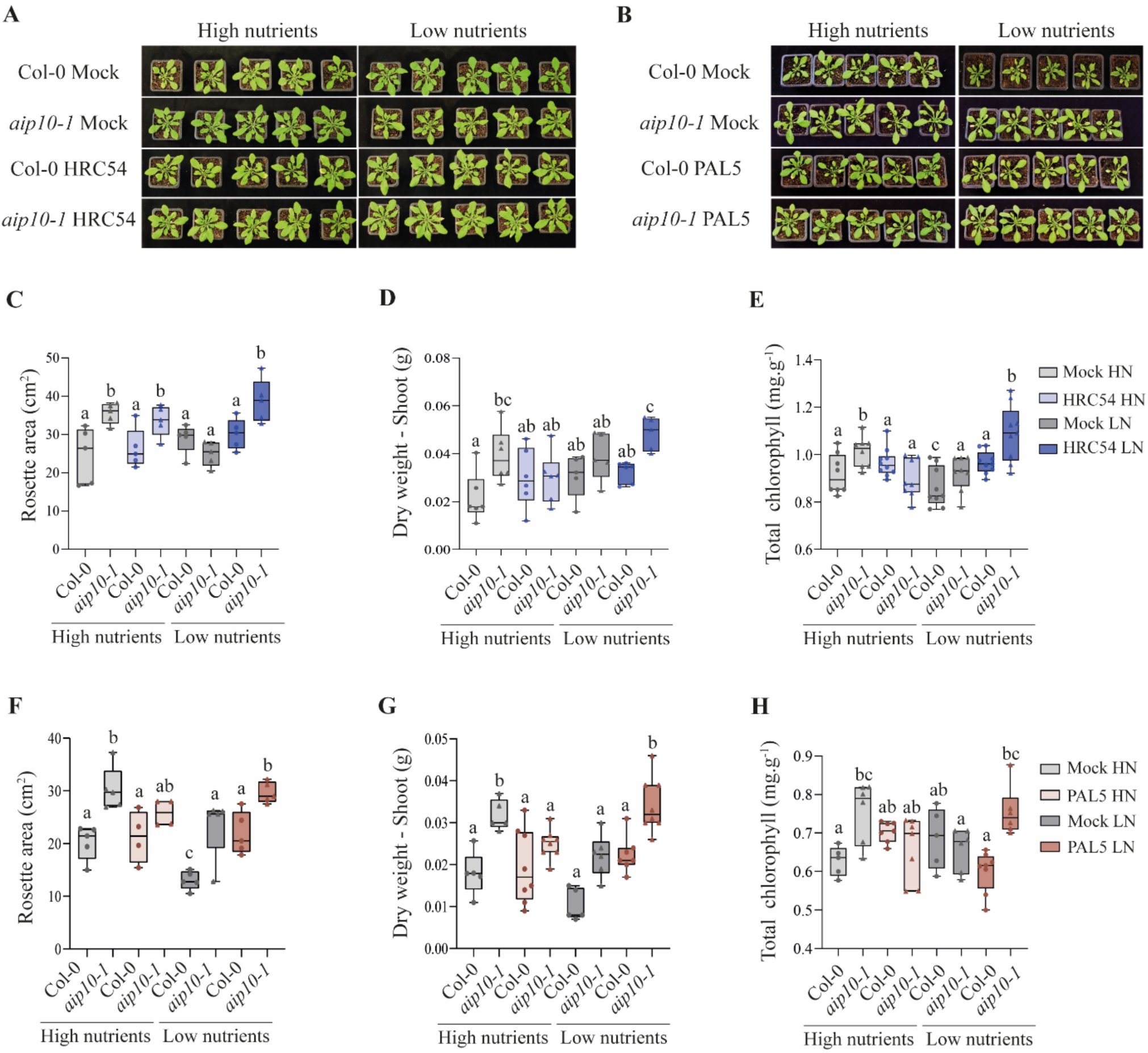
Shoot development of *aip10-1* and wild-type Col-0 plants inoculated with different plant growth-promotion bacteria, grown in a greenhouse under low and high fertilization conditions (0.06 g/L and 0.12 g/L NPK, respectively). (A) Representative images of *A. thaliana* Col-0 and *aip10-1* plants inoculated with *H. seropedicae* HRC54 at 35 DAG (31 DAI), grown in a greenhouse under two different fertilization conditions. **(B)** Representative images of *A. thaliana* Col-0 and *aip10-1* plants inoculated with *G. diazotrophicus* Pal5 at 35 DAG (31 DAI), grown in a greenhouse under two different fertilization conditions. **(C to E)** mock and HRC54-inoculated plants at 35 DAG. **(C)** Rosette area measurement using ImageJ program; **(D)** Shoot dry weight; **(E)** Total chlorophyll content in leaf disks. **(F to H)** mock and Pal5-inoculated plants at 35 DAG. **(F)** Rosette area measurement using ImageJ program; **(G)** Shoot dry weight; **(H)** Total chlorophyll content in leaf disks. The box plot graphs show the distribution of data of six individuals. The box represents the interquartile range (IQR), with the inner line running down the median. The whiskers extend to the maximum and minimum values, including all points within that range. Differences between groups were confirmed by Student’s t-test and means followed by different letters indicate significant differences at the 5% confidence level

Root phenotypic analyses showed that *aip10-1* plants exhibited more developed root systems in response to inoculation with the HRC54 and Pal5 strains upon low fertilization (Fig. 6). Under these conditions, *aip10-1* plants inoculated with HRC54 and Pal5 exhibited 20–51% longer roots (Fig. 6B and E) and 24–42% larger surface area (Fig. 6C and F) compared to Col-0, respectively. At the same time, under high fertilization, non-inoculated *aip10-1* plants still exhibited average increases of 40% in root length and 50% in surface area relative to wild-type plants (Fig. 6B-C and 6E-F). The increased root growth of *aip10-1* mutants resulted in a significant increase in dry biomass. In response to HRC54 and Pal5 inoculation, the gain in root dry weight ranged from 40% (Fig. 6A) to 70% (Fig. 6D) in mutant plants compared to wild-type plants under low fertilization, equivalent to the increase found in *aip10-1* mock-treated plants grown under high fertilization. Under low-nutrient conditions, Biofree inoculation markedly enhanced root biomass, elongation, and surface area in mutant plants, whereas no significant effects were observed in Col-0 (Fig. S5).

**Figure 6.**
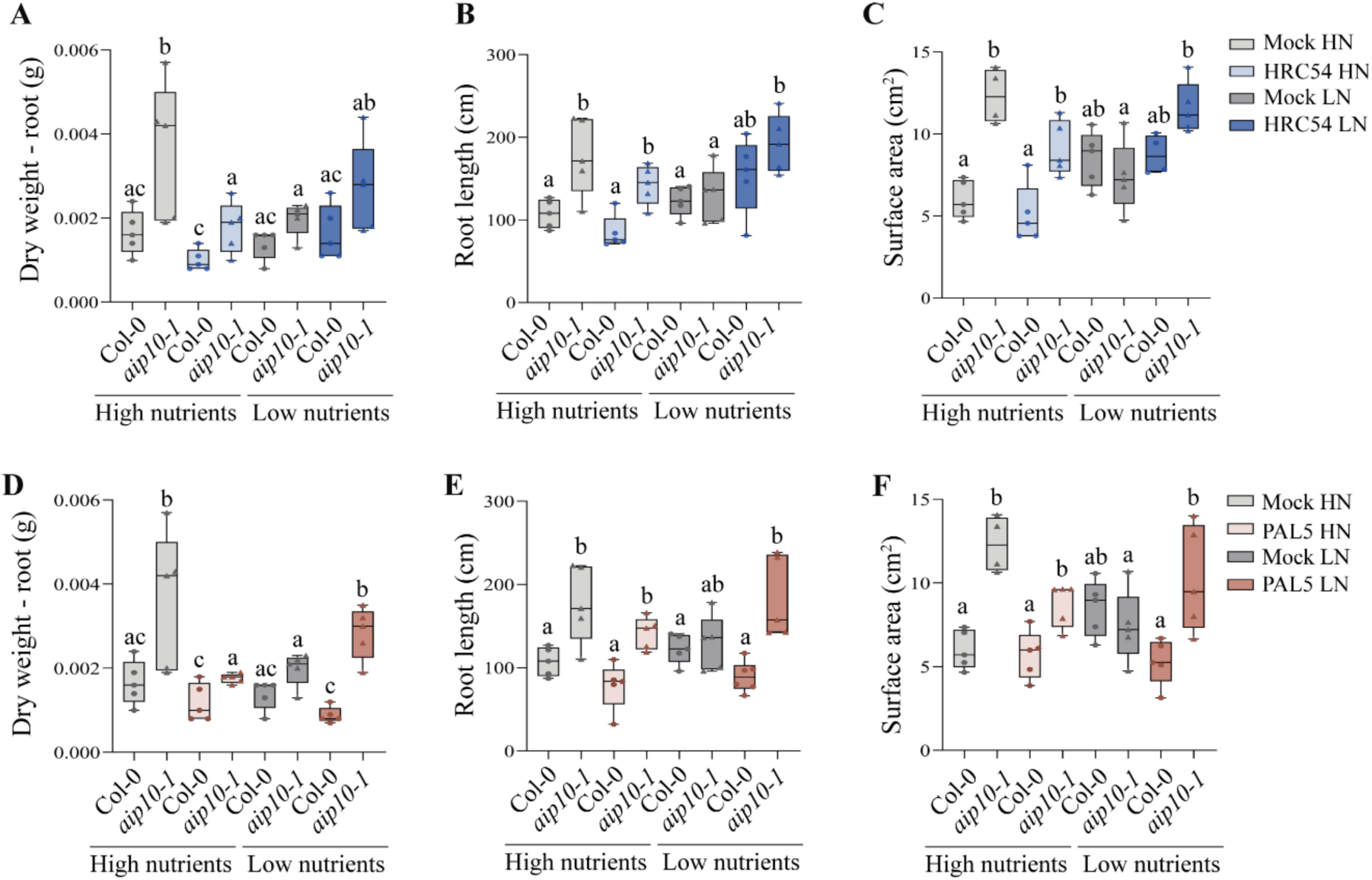
Root development of Col-0 and *aip10-1* plants inoculated with different plant growth-promoting bacteria, grown in a greenhouse under low and high fertilization conditions at 35 DAG. (A) Root dry weight in mock and HRC54-inoculated plants. **(B)** Root length in mock and HRC54-inoculated plants. **(C)** Root surface area in mock and HRC54-inoculated plants. **(D)** Root dry weight in mock and Pal5-inoculated plants. **(E)** Root length in mock and Pal5-inoculated plants. **(F)** Root surface area in mock and Pal5-inoculated plants. Root length and surface area were measured by WinRHIZO. The box plot graphs show the distribution of data of five individuals. The box represents the interquartile range (IQR), with the inner line running down the median. The whiskers extend to the maximum and minimum values, including all points within that range. Differences between groups were confirmed by Student’s t-test and means followed by different letters indicate significant differences at the 5% confidence level.

The use of bioinoculants also promoted growth at the plant reproductive stage (Fig. 7A). Consistent with the results for the vegetative phase, low fertilization positively influenced the height of the main inflorescence axis in inoculated mutant plants, in response to both inoculations, compared to Col-0 (Fig. 7B and E). In addition, this combination of stimuli generated inflorescences with 23–29% more siliques on the main axis of inoculated *aip10-1* plants compared to all other groups grown under low fertilization (Fig. 7C and F). Similarly, *aip10-1* mutants produced a higher total seed weight under low fertilization conditions, following inoculation of HRC54 (Fig. 7D) and Pal5 (Fig. 7G). Additionally, the commercial microbial consortium more strongly increased silique number and total seed yield in *aip10-1* than in Col-0, particularly under low nutrient conditions (Fig. S6). Overall, *aip10-1* mutants responded more strongly to bioinoculants containing diazotrophic bacteria, resulting in greater dry biomass accumulation in both shoots and roots compared to Col-0 plants. Moreover, these benefits were primarily evident under low fertilization conditions. Taken together, the results suggest that PGPB bioinoculant application can partially replace mineral fertilization in *aip10-1* mutant plants, but not in Col-0, highlighting a genotype-specific response.

**Figure 7.**
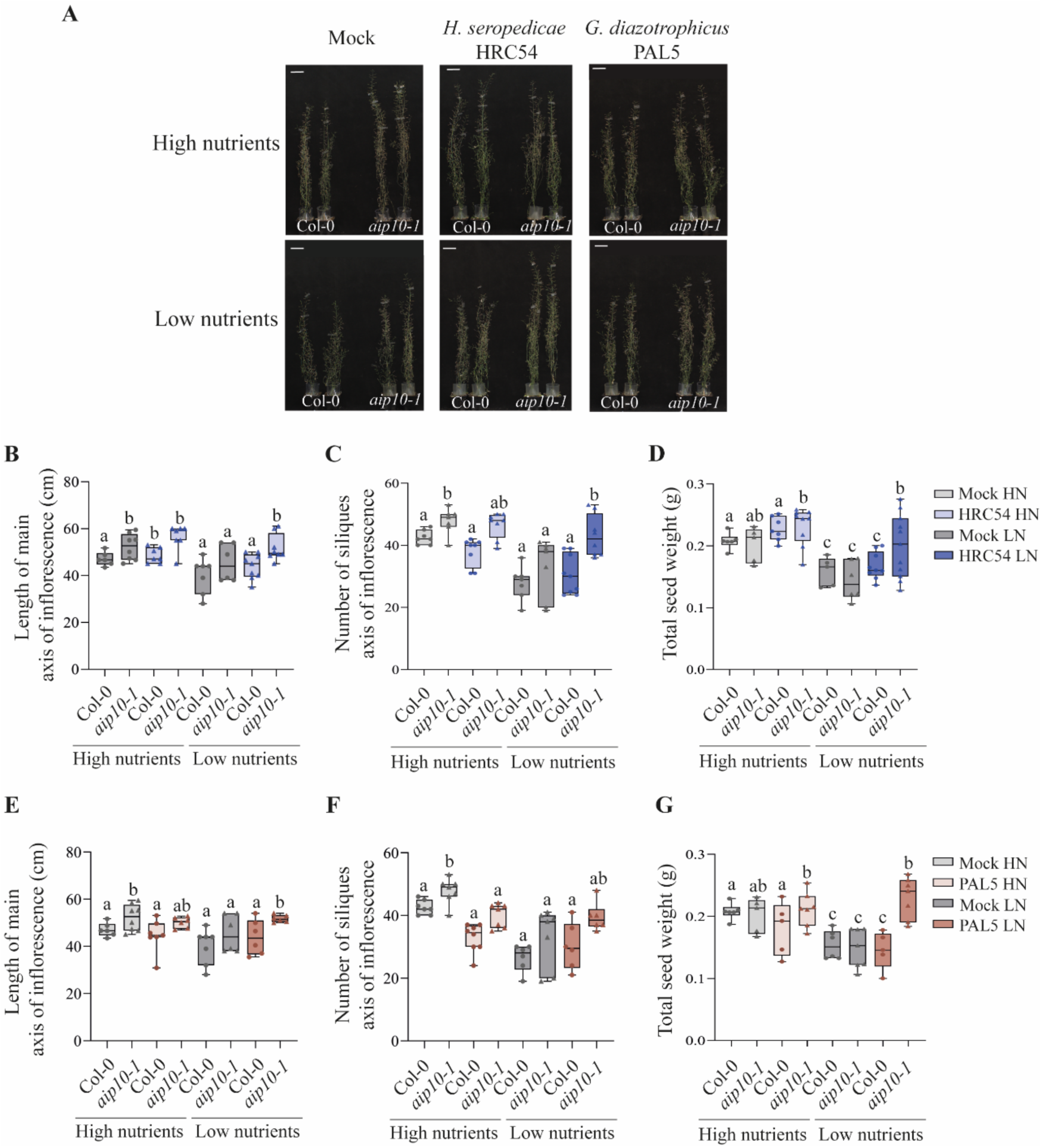
Reproductive stage of Col-0 and *aip10-1* plants inoculated with different plant growth-promoting bacteria, cultivated in a greenhouse under low and high fertilization conditions at 60 DAG. (A) Representative images of Col-0 and *aip10-1* during the reproductive stage comparing the different responses to HRC54 and Pal5 inoculation, in response to low and high fertilization treatments. **(B to D)** mock and HRC54-inoculated plants; **(B)** Length of the main inflorescence axis; **(C)** Total number of siliques in the main inflorescence axis; **(D)** Total seed weight per individual. **(E to G)** mock and Pal5-inoculated plants; **(E)** Length of the main inflorescence axis; **(F)** Total number of siliques in the main inflorescence axis; **(G)** Total seed weight per individual. The box plot graphs show the distribution of data of five individuals. The box represents the interquartile range (IQR), with the inner line running down the median. The whiskers extend to the maximum and minimum values, including all points within that range. Differences between groups were confirmed by Student’s t-test and means followed by different letters indicate significant differences at the 5% confidence level.

### Nitrogen metabolism in *aip10-1* plants associated with diazotrophic PGPB under high and low fertilization conditions

Our data showed that PGPB inoculation in *aip10-1* increased bacterial *nifH* expression and had additive effects on plant biomass and chlorophyl levels. We therefore investigated N assimilation in *aip10-1* plants by analyzing the expression of representatives in two selected key gene families known to respond to changes in N availability and PGPB inoculation (Rosman *et al*., 2024). The N assimilation genes analyzed were: i) the two Nitrate Reductases (NR) *NIA1* and *NIA2* that are modulated by nitrate availability (Wilkinson and Crawford, 1993); ii) and the two Glutamine Synthetase cytosolic genes (GS1), *GLN1;1* and *GLN1;3,* that are high-affinity and low-affinity-related isoforms, respectively (Ishiyama *et al*., 2004). Analyzes were performed by RT-qPCR of root samples collected at 20 DAG from wild-type and *aip10-1* plants mock-treated or PGPB inoculated and grown in a greenhouse under the high and low fertilization conditions.

We first evaluated the N metabolism data in mock-treated plants, as the native microbiome in non-inoculated *aip10-1* root compartment was enriched with PGPB and diazotrophic bacteria. The effects of nutrient availability on *nifH* expression were evaluated in natural microbiomes associated with wild-type and *aip10-1* roots. Consistent with the microbiome data enriched with diazotrophic PGPB, higher *nifH* expression levels were observed in mock-treated *aip10-1* plants grown under low nutrient availability (Fig. 8A). Analysis of nitrogen assimilation pathways showed that *NIA1* and *NIA2* were induced under high nutrients availability in mock-treated *aip10-1* plants (Fig. 8B and C), as was *GLN1;3* (Fig. 8E), compared to wild-type Col-0. No modulation of these genes was observed under low nutrients condition. Under both nutritional conditions, mock-treated *aip10-1* plants showed a tendency towards higher levels of *GLN1;1* expression compared to Col-0 (Fig. 8D).

**Figure 8.**
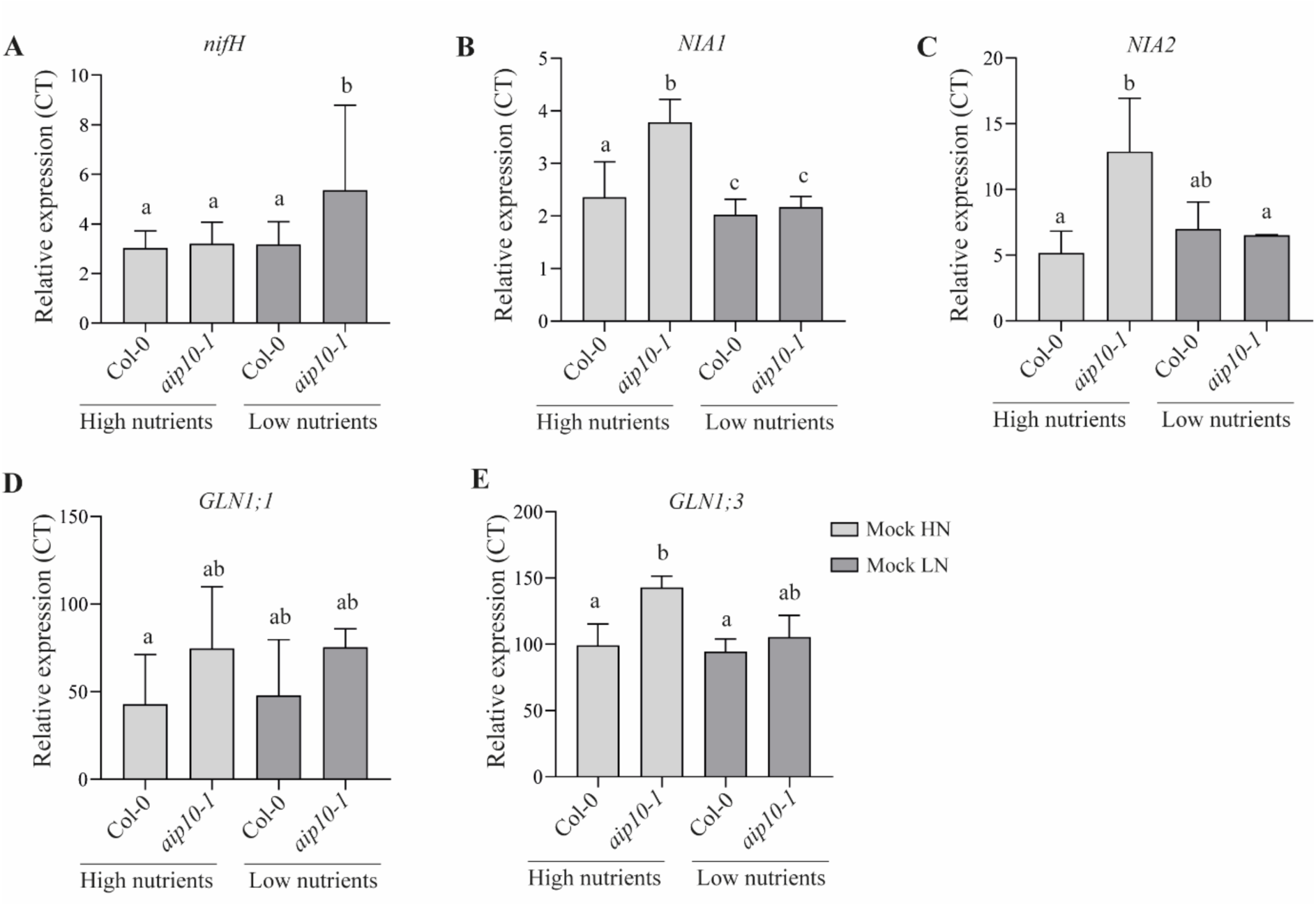
Expression of nitrogen metabolism-related genes in mock-treated Col-0 and *aip10-1* roots, cultivated in greenhouse under high and low fertilization conditions at 20 DAG. qRT-PCR analyses were performed in mock-treated Col-0 and *aip10-1* roots, cultivated under high and low fertilization conditions at 20 DAG. (**A**) *nifH* transcript abundance. (B) *NIA1* transcript levels. (C) *NIA2* transcript levels. (**D**) *GLN1;1* transcript levels. (**E**) *GLN1;3* transcript levels. Relative expression values were quantified by RT-qPCR and normalized by *UBI14* and *GAPDH* reference genes. Bars represent mean ± standard deviation of the relative mRNA expression in three biological replicates and each biological replicate analyzed with three technical replicates. Statistical analysis was performed using the Student’s t-test (P ≤ 0.05). Means with different letters are significantly different at 5% level of confidence. This mock-treated Col-0 and *aip10-1* expression data is the same as the one presented in Fig.9.

Taken together, the data in mock-treated plants showed that the use of high concentration of mineral fertilizers negatively affected the diazotrophic PGPB root microbiome in *aip10-1* roots. However, under these conditions, *AIP10* silencing could activate ammonium and nitrate assimilation pathways, suggesting that *aip10-1* assimilates nitrogen more efficiently compared to wild-type plants. This is consistent with the enhanced biomass accumulation previously reported for *aip10-1* compared to Col-0, growing under high nutrient conditions, further supporting the genotype-specific response.

We next evaluated N metabolism in PGPB inoculated plants, as diazotrophic PGPB inoculation of *aip10-1* under low nutrient availability enhanced growth promotion. The control mock-treated Col-0 and *aip10-1* expression data is the same as the one presented in Fig.8. The effects of nutrient availability on *nifH* expression showed that low nutrient availability increased *nifH* expression in *aip10-1* plants inoculated with HRC54 and Pal5 relative to high-nutrient conditions or mock-inoculated controls (Fig. 9A and B). Under low N conditions, *GLN1;1* expression in inoculated *aip10-1* plants showed a tendency toward higher levels, although no significant differences were detected relative to mock controls or to mutant plants grown under high nutrient availability (Fig. 9C and D). In contrast, under low N fertilization, *H. seropedicae* inoculation significantly increased *NIA1* expression in *aip10-1* plants compared to other inoculated groups grown under high or low fertilization, whereas *G. diazotrophicus* inoculation had no effect on *NIA1* expression (Fig. 9E and F). Altogether, these findings suggest that although *nifH* expression was increased in diazotrophic PGPB inoculated *aip10-1* plants, the observed growth promotion may not be explained exclusively by biological N₂ fixation. Additional mechanisms may contribute, including PGPB-driven increases in nutrient solubilization and mobilization, nutrient use efficiency (NUE), as well as shifts in the composition of the root-associated microbial community.

**Figure 9.**
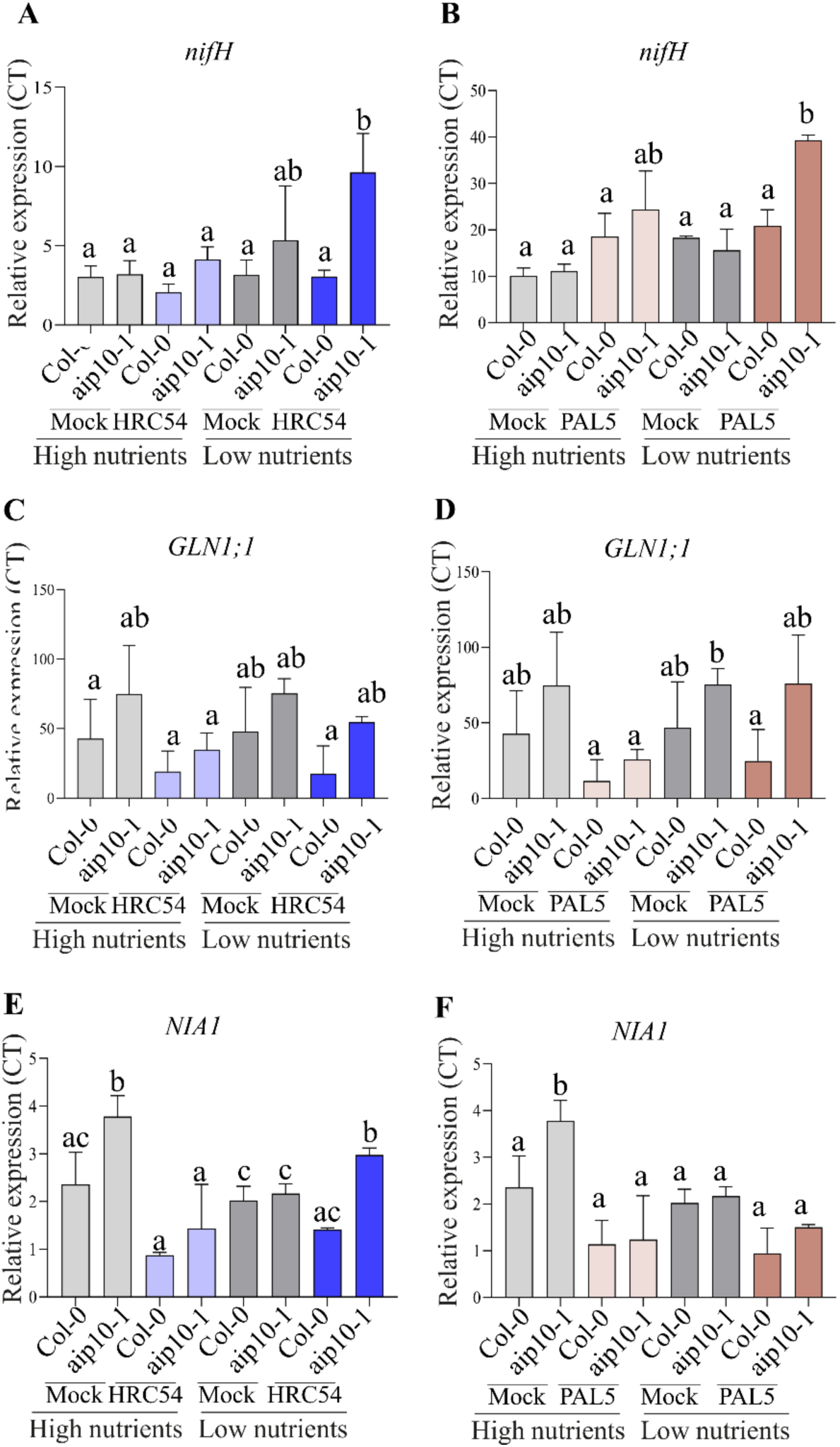
Expression of nitrogen metabolism-related genes in Col-0 and *aip10-1* roots of inoculated plants, cultivated in greenhouse under high and low fertilization conditions at 20 DAG. (A, B) *nifH* transcript abundance from *H. seropedicae* HRC54 and *G. diazotrophicus* Pal5, respectively, colonizing Col-0 and *aip10-1* roots. (C, D) *GLN1;1* mRNA level in *H. seropedicae* HRC54 and *G. diazotrophicu*s Pal5 inoculated roots, respectively. (E, F) *NIA1* transcript levels in *H. seropedicae* HRC54 and *G. diazotrophicu*s Pal5 inoculated roots, respectively. Relative expression values were quantified by RT-qPCR and normalized by UBI14 and GAPDH reference genes. Bars represent mean ± standard deviation of the relative mRNA expression in three biological replicates and each biological replicate analyzed with three technical replicates. Statistical analysis was performed using the Student’s t-test. Means with different letters are significantly different at 5% level of confidence. This mock-treated Col-0 and *aip10-1* expression data is the same as the one presented in Fig.8.

## 4. Discussion

Achieving high agricultural productivity while minimizing environmental harm requires the adoption of novel and sustainable approaches that limit dependence on mineral fertilizers. PGPB bioinoculants represent a promising strategy to enhance crop performance while reducing reliance on mineral fertilizers. However, their effectiveness is highly variable and largely depends on plant genetic factors that regulate microbial recruitment and colonization. Although plants actively shape their root-associated microbiota, the molecular mechanisms that enable the accommodation of beneficial PGPB without triggering defense responses, culminating with the promotion of plant growth and yield, remain poorly understood (Vandenkoornhuyse *et al*., 2015; Zhang *et al*., 2023). Elucidating these plant-controlled processes is critical for improving the consistency of PGPB-plant interactions.

Our research group has systematically investigated the molecular basis of plant-PGPB communication, identifying key plant signaling, transcriptional, and metabolic pathways that are modulated during these interactions ultimately influencing plant physiological and developmental responses (Carvalho *et al*., 2011, 2016, 2022; Thiebaut *et al*., 2017; Hardoim *et al*., 2020; Ballesteros *et al*., 2021; Rosman *et al*., 2024). Here, we show that among the sugarcane gene networks regulated by inoculation with PGPB is a regulator of cell cycle progression that may modulate plant growth by integrating cell divisions to abiotic and biotic environmental signals. Within this network, AIP10 integrates cell division rates with metabolic regulation, by direct interaction with ABAP, SnRK1 (via KIN10) and possibly additional partners. AIP10 acts as a negative regulator of the cell cycle, and its silencing results in healthier plants with increased biomass and yield, along with enhanced metabolite accumulation, including carbohydrate content (Montessoro *et al*., 2025). Therefore, we investigated whether *AIP10* silencing contributes to beneficial interactions with PGPB in the endogenous root microbiome or in response do PGPB-bioinoculants, by performing functional analyses in *A. thaliana aip10-1* knockout mutant. Our results showed that the improved performance of *A. thaliana aip10-1* was associated with greater colonization by beneficial bacteria, both within the native root microbiome or following PGPB inoculation. Based on these findings and recent advances in plant–PGPB interactions, we propose potential mechanisms that may explain the enhanced PGPB colonization in *aip10-1*, contributing to the observed increase in plant performance (Figure 10).

**Figure 10.**
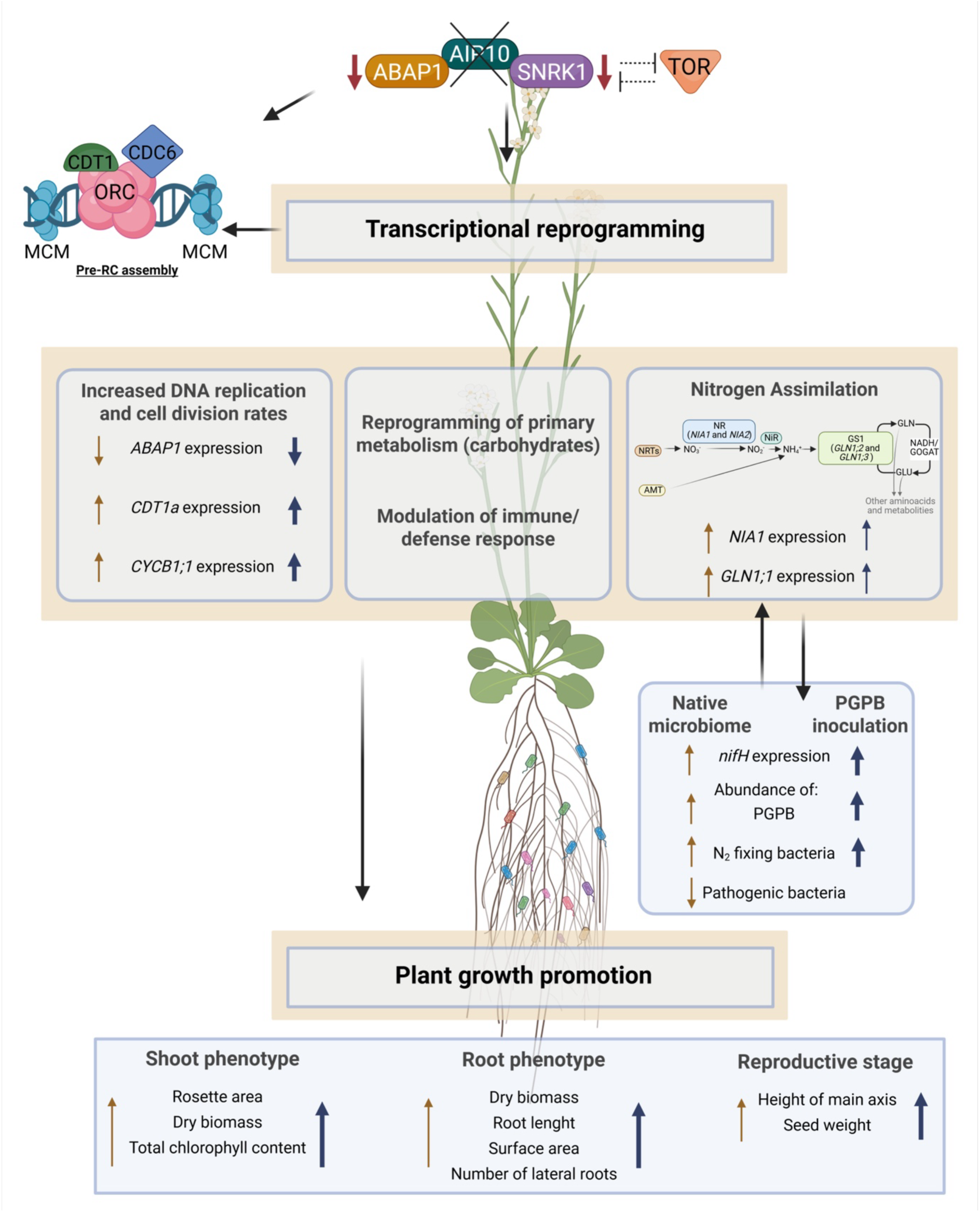
A hypothetical model illustrating the potential mechanisms that coordinate plant growth and metabolism with the recruitment of beneficial root microbiota in the *AIP10* knockout mutant of *A. thaliana,* under low nutrient conditions. AIP10 is proposed to function as a regulatory node connecting cell cycle control and primary metabolism, likely through interactions with ABAP1, SnRK1 (via KIN10), and possibly additional partners. Elimination of AIP10 in *aip10-1* mutant triggers broad transcriptional changes across multiple pathways. Attenuation of AIP10/ABAP1 signaling enhances pre-replication complex (Pre-RC) assembly by relieving repression of pre-RC genes (*CDT1a)*, resulting in increased DNA replication and higher cell division rates (*CYCB1;1*). This process is accompanied by transcriptional reprogramming of metabolic pathways leading to greater carbohydrate accumulation, along with modulation of immune and defense responses, and nitrogen assimilation genes (*NIA1* and *GLN1;1*), in a manner that favors the association with beneficial plant growth promoting bacteria. At the root level, non-inoculated *aip10-1* plants exhibit enrichment of beneficial microbial taxa, including diazotrophic bacteria, and a reduced abundance of pathogenic taxa, comparing to Col-0. Under low nitrogen conditions, PGPB inoculation further enhances bacterial colonization and further induces *nifH* expression in *aip10-1* compared to Col-0. The molecular and microbiome shifts promote improved root system architecture, increased shoot biomass and chlorophyll content, and enhanced reproductive performance, with the strongest growth responses observed in inoculated *aip10-1* plants under low nitrogen availability. Orange arrows indicate responses in mock-treated *aip10-1* plants compared to Col-0, whereas blue arrows represent responses in inoculated *aip10-1* plants compared to Col-0 plants. Arrow thickness corresponds to the magnitude of the response. Created with BioRender.com.

### *AIP10* silencing favored beneficial microbial recruitment

Increasing evidence suggests that plant performance and environmental adaptation rely on coordinated interactions with the root microbiota. Data are emerging showing that genes governing development, immune responses, nutrient uptake, and metabolite release can possibly function as determinants of the microbiome assembly (Zhang *et al*., 2023). Consistently, we found that the loss of *AIP10* function in *A. thaliana* reshaped root-associated bacterial communities while maintaining a substantial core of shared taxa. Notably, *aip10-1* plants preferentially recruited beneficial PGPB, particularly diazotrophic species, in both the root compartment and rhizospheric soil, while being depleted in potentially pathogenic taxa.

The mechanisms by which *aip10-1* re-shapes its root microbiome may involve alterations in plant carbohydrate metabolism, reported as key nutrient sources for PGPB (Upadhyay *et al*., 2022; Hemelda and Noutoshi, 2025). Carbohydrate levels were quantitatively increased in *aip10-1* plants indicating changes in primary metabolism (Montessoro *et al*., 2025). Consistently, genes associated with carbohydrate metabolic pathways were found as differentially expressed in *aip10* roots, suggesting that altered carbon availability could support the recruitment of diazotrophic PGPB. Root-derived metabolites were also shown to play an actively selective role in shaping the root microbiome, as mutant in genes involved in regulating specialized metabolites influenced microbial recruitment, community structure, and function (Vandenkoornhuyse *et al*., 2015; Zhang *et al*., 2023). Loss-of-function mutations in genes controlling coumarin biosynthesis or export significantly modified rhizosphere and endosphere communities, promoting the recruitment of growth-promoting or nutrient-mobilizing bacteria through changes in root-derived specialized metabolites (Stringlis *et al*., 2018; Huang *et al*., 2019; Voges *et al*., 2019). Secondary metabolites such as phenolics and terpenes have been shown to modulate microbiome composition and enrich beneficial taxa (Stringlis *et al*., 2018; Harbort *et al*., 2020). Furthermore, genotype-specific metabolites explain substantial microbiome variation in crops such as rice (Liu *et al*., 2025).

In parallel, in rice, NRT1.1B, a nitrate transporter and sensor, was associated with modulation of root microbiota composition, integrated with higher NUE in field-grown plants (Zhang *et al*., 2019). Consistent with this mechanism, non-inoculated *aip10-1* plants showed higher expression of N assimilation genes compared to Col-0, along with a root microbiota enriched in PGPB and diazotrophic bacteria that enhance N availability to the plant.

*aip10-1* mutants showed transcriptional reprogramming of genes in pathways related to plant-microorganism interaction such as cell wall formation, hormonal responses, defense and immune response to bacteria. In *aip10-1* roots, induction of genes involved in cell wall loosening, such as *XTH* and *PME46*, and repression of genes associated with cell wall reinforcement and stress-related remodeling, including *EXT15* and *LRX3,* suggest increased cell wall plasticity. This shift may facilitate bacterial association and root tissue colonization. In sugarcane, genotypes that efficiently associate with endophytic diazotrophic bacteria show extensive reprogramming of cell wall–modifying genes, promoting increased tissue plasticity and microbial accommodation (Ballesteros *et al*., 2021). Consistent with this, broader evidence indicates that cell wall remodeling is a key feature of plant readiness for symbiotic or associative interactions, as exposure to PGPR commonly induces expansins and XTHs across diverse plant species (Vacheron *et al*., 2013; Srivastava *et al*., 2023).

Alterations in innate immunity pathways may also lead to distinctive microbial assemblages, as shown in mutants impaired in pattern-recognition receptors, which exhibit shifts in core bacterial taxa and increased abundance of taxa typically associated with beneficial functions (Ma *et al*., 2021). In *aip10-1* roots, some genes involved in immune and defense pathways related to bacterial interactions were either repressed or induced. Such transcriptional reprogramming is required for successful beneficial associations with PGPB, as plants must downregulate specific defense responses to permit bacterial colonization while simultaneously activating other defense pathways to prevent bacterial overgrowth (Carvalho *et al*., 2016). Furthermore, it might participate in the regulation how roots perceive and respond to soil microorganisms, creating a more permissive environment for colonization by diazotrophic and PGPB-enriched taxa while limiting the presence of potential pathogens. Upregulation of bacterial response regulators such as ILA in *aip10-1* is consistent with plants genetically primed for beneficial microbial associations showing enhanced sensitivity to microbe-derived signals before colonization (Liu *et al*., 2017; Gao *et al*., 2022).

The transcriptional profile of *aip10-1* plants in the absence of PGPB inoculation resembles signatures reported in plant genotypes with enhanced capacity to interact with beneficial microorganisms. It suggests that *AIP10* silencing may induce a primed physiological state that facilitates the establishment of growth-promoting associations with PGPB.

### Enhanced growth promotion in *aip10-1* in response to PGPB inoculation were driven by cell division and nutrient availability

Our data suggest that mock *aip10-1* genotype is primed for enhanced association with PGPB bioinoculants. This genotype appears to provide physiological conditions that favor colonization by beneficial and diazotrophic bacteria, thereby increasing plant’s capacity to respond to their growth-promoting effects. Consistent with a native root microbiota enriched with beneficial bacteria, inoculated *aip10-1* genotype was colonized by higher numbers of PGPB bioinoculant during early phases after inoculation. Consequently, *aip10-1* plants exhibited increased root and shoot dry biomass and higher productivity in response to inoculation with distinct PGPB, such as *G. diazotrophicus* Pal5, *H. seropedicae* HRC54, and Biofree, compared to inoculated wild-type plants. Moreover, the enhanced traits observed in non-inoculated *aip10-1* plants were further amplified upon inoculation, indicating additive effects on plant growth promotion.

Increased plant dry biomass ultimately depends on higher rates of cell division, as plant growth and morphogenesis are governed by the balance between cell division, cell expansion, and cell differentiation (Hemerly et al. 1999, Jorgensen and Tyers, 2004; Jones *et al*., 2017; Carneiro *et al*., 2021). AIP10 is part of a regulatory network that controls plant growth by modulating cell cycle progression at G1 to S phase, a key checkpoint regulated by environmental signals, such as biotic interactions (Masuda *et al*., 2008; Montessoro *et al*., 2025). ABAP1 and AIP10 negatively regulate DNA replication by reducing the activity of the pre-replication complex (preRC) and by decreasing transcription of preRC members as *AtCdt1a* and *AtCdt1b* (Masuda *et al*., 2008; Montessoro *et al.,* 2025). In addition, *ABAP1* expression is reduced in *aip10-1* plants suggesting that AIP10 may also control cell division through a feedback mechanism of *ABAP1* transcriptional control. Our results suggest that the enhanced growth of *aip10-1* plants in response to bioinoculants can be increased by the negative regulation of *ABAP1*, favoring higher cell division rates and increased growth. Moreover, *AtCDT1a* showed increased expression in *aip10-1* plants after inoculation with both bioinoculants. A similar pattern was observed for the cell division marker *CYCB1;1*, supporting the idea that beneficial bacterial colonization could stimulate higher cell division rates in *aip10-1*.

PGPB associations can promote plant growth and increase dry biomass through mechanisms that enhance nutrient availability such as biological nitrogen fixation and mineral solubilization, and by improvement of nutrient use efficiency (Thiebaut *et al*., 2022). Here, we showed that nutrient availability modulated the interaction between PGPB and *aip10-1* plants, with an intensified growth-promoting response under low-fertilization conditions. PGPB inoculation in low nutrient availability resulted in enhanced growth similar to that observed in non-inoculated mutant plants grown under high fertilization (Montessoro *et al*., 2025). Under these conditions, inoculation increased shoot biomass accumulation by up to 35% in *aip10-1* plants under low fertilization. Silencing *AIP10* promoted the formation of more robust root systems in response to inoculation with HRC54 and Pal5 under low fertilization, increasing root surface area by up to 42% compared to Col-0. This enhanced root growth resulted in a significant increase in dry biomass, with gains of up to 70% after inoculation, equivalent to the increase observed in non-inoculated *aip10-1* plants grown under high fertilization.

N levels in the environment regulate the expression of multiple genes involved in assimilation, transport, and utilization (Wilkinson and Crawford, 1993; Lothier *et al*., 2011; O’Brien *et al*., 2016). In this study, we investigated the expression pattern of members of GS and NR gene families, which regulate N assimilation and are modulated by variations in N availability, particularly in the presence of PGPB (Rosman *et al*., 2024). High nutrient availability induced the expression of *NIA1*, *NIA2*, and *GLN1;3*, in mock-treated aip*10-1* plants compared to wild-type plants, with no additional effect of inoculation. Under low fertilization conditions, *NIA1* was induced in *H. seropedicae* inoculated *aip10-1.* Transcriptome analysis revealed that *NIA1* was induced in shoots and roots of *A. thaliana* in response to inoculation with Pal5, at 50 DAI (Soares *et al*., 2024). Moreover, inoculation with the endophytic fungus *Piriformospora indica* induced NR gene expression, resulting in increased N accumulation in *A. thaliana* and tobacco seedlings (Sherameti *et al*., 2005).

From the bacterial perspective, *nifH* expression in HRC54 and PAL5 inoculated *aip10-1* plants was consistently higher compared to Col-0 for both strains at 7 DAI. Remarkably, this difference persisted at 16 DAI only under low-nutrient conditions. Given that the plant growth-promoting capacity of *H. seropedicae* and *G.* relies on coordinated N₂-fixation–related gene activity (Pedrosa *et al*., 2011; Tufail *et al*., 2021), our results suggest that *AIP10* loss of function of *AIP10* not only enhances root colonization by PGPB, but also potentiates bacterial N_2_-fixation metabolism, especially in environments with low N availability.

The efficiency of the interaction between plants and PGPB has been widely attributed to external conditions, including plant genotype, bacterial genotype, and soil nutrient availability (Ballesteros *et al*., 2021; Carvalho *et al*., 2022; Rosman *et al*., 2024). Among these factors, the nutritional composition of the soil stands out, playing an essential role in controlling plant growth and development, including the establishment of the plant-PGPB association. Low N fertilization conditions encourage the establishment of efficient plant-bacteria associations (Tufail *et al*., 2021; Rosman *et al*., 2024).

In sugarcane, the association with PGPB can contribute up to 70% of all accumulated N through BNF or improved NUE, varying according to the plant and bacterial genotype and soil fertility (Urquiaga *et al*., 2012; Antunes *et al*., 2019; Martins *et al*., 2020; Singh *et al*., 2021). Among the diazotrophic bacteria that are associated with sugarcane, the *Herbaspirillum* stands out, found colonizing the apoplast of leaves, very close to the vascular parenchyma cells (Silva *et al*., 2003). Several studies have already reported the positive effects of this association, such as promoting increased biomass, shoot and secondary root growth, as well as the accumulation of N, P and K (Oliveira *et al*., 2006; dos Santos *et al*., 2020; Martins *et al*., 2020). Specifically for *H. seropedicae* in maize, inoculation has been shown to increase root fresh mass, root surface area (Nunes *et al*., 2021); number of lateral roots (Do Amaral *et al*., 2014; Ferrari *et al*., 2014; Hardoim *et al*., 2020); N accumulation (Brusamarello-Santos *et al*., 2017); in addition to altering N metabolism and ensuring tolerance to abiotic stresses (Alves *et al*., 2014; Curá *et al*., 2017; Breda *et al*., 2019; Dias *et al*., 2021). Moreover, *aip10-1* plants inoculated with Biofree showed increased rosette expansion and biomass accumulation compared with mock-treated mutants under low nutrient availability. As this commercial bioinoculant comprises *A. brasilense* Ab-V6 and *P. fluorescens* CCTB03, PGPB associated with phytohormone synthesis and phosphate solubilization (Park *et al*., 2009; Zhao *et al*., 2025), these results suggest that multiple growth-promoting mechanisms may contribute to enhanced plant performance in plants with silenced *AIP10* expression under reduced chemical fertilization.

### AIP10 as a regulatory target for improving plant performance under nutrient-limited conditions in association with PGPB

Conventional strategies for maintaining high crop productivity largely depend on the extensive application of nitrogen-based fertilizers. However, crop yields have not increased proportionally to fertilizer inputs, revealing low NUE and higher production costs (Mueller *et al*., 2019). In this context, to meet the nutritional needs of crops and maintain high levels of productivity, modern agriculture has resorted to the use of bio-inputs, which favor plant production, soil health, and the reduction of GHG emissions. Therefore, investing in alternatives such as modulation of native root microbiomes to favor enrichment with PGPB, or development PGPB-based biological inputs, becomes crucial strategies to ensure the agricultural sustainability of various crops such as maize, sugarcane, wheat, and rice (Canellas *et al*., 2013; Martins *et al*., 2020; Carril *et al*., 2021; Pankievicz *et al*., 2021; Tufail *et al*., 2021). Dissecting the plant genetic factors that govern the attraction of beneficial PGPB within the microbiome, as well as the establishment of effective interactions with PGPB bioinoculants, is essential for improving crop yield and plant fitness.

The absence of AIP10 shaped root microbiome to a higher abundance of PGPB and diazotrophic bacteria. This technology also increased responsiveness to colonization by bioinoculants under low nutrient availability. Remarkably, the enhanced traits observed in non-inoculated *aip10-1* plants were further amplified upon inoculation, indicating additive effects on plant growth promotion. Our observations raise the important question of why the loss of AIP10 function, a gene conserved across terrestrial plants, confer advantages such as enhanced root colonization by beneficial bacteria. Loss of AIP10 activity confers traits associated with enhanced terrestrial resilience, including increased CO₂ assimilation, reduced transpiration, and improved water-use efficiency. We therefore hypothesize that AIP10 contributes to the regulation of soil–microbe interactions as part of plant adaptation to terrestrial environments.

Here, we investigated the modulation of the native root microbiome and the response to bioinoculant inoculation in *A. thaliana aip10-1* plants, aiming to enhance plant growth and biomass accumulation under reduced chemical fertilization. Under low fertilization conditions, plants lacking AIP10 shaped their root microbiome toward a higher abundance of PGPB and diazotrophic bacteria and displayed improved responsiveness to bioinoculants, accumulating biomass comparable to that of non-inoculated plants grown under high fertilization. In addition, similarly to the vegetative phase, low fertilization increased reproductive development, leading to a 29% increase in siliques on the main inflorescence axis compared with all other groups under the same conditions. Taken together, our results indicate that modifying *AIP10* expression, previously associated with the control of a central hub that integrates cell division rates and primary metabolism, can be used as a strategy to improve nutrient use efficiency under low chemical fertilization, contributing to a more sustainable and regenerative agriculture. This approach results in a real increase in biomass, productivity, and possibly the nutritional value of the plants, having an important impact on food security.

## Supplementary data

The following supplementary data are available at JWB online.

Supplementary Table S1. Primers used in this study.

Supplementary Table S2. Relative abundance of diazotrophic, plant growth–promoting (PGPB), and pathogenic bacterial communities associated with the roots of Col-0 and *aip10-1* plants.

Supplementary Fig. S1. Differential gene expression patterns in Col-0 and *aip10-1* roots of *A. thaliana* 11 days after germination.

Supplementary Fig S2. Visualization of bacterial colonization patterns in different root zones of *A. thaliana* Col-0 and *aip10-1* mutant plants.

Supplementary Fig S3. Experimental design of the differential fertilization trial in *aip10-1* and wild-type Col-0 plants inoculated with *H. seropedicae* HRC54 (HRC54), *G. diazotrophicus* Pal5 (Pal5), Biofree or non-inoculated (mock).

Supplementary Fig S4. Growth performance and physiological responses of Col-0 and *aip10-1* plants inoculated with commercial bioinoculant Biofree under contrasting nutrient conditions.

Supplementary Fig S5. Root architecture responses of Col-0 and *aip10-1 A. thaliana* plants to Biofree inoculation under contrasting nutrient conditions.

Supplementary Fig S6. Reproductive performance of Col-0 and *aip10-1 A. thaliana* plants in response to Biofree inoculation under high- and low-nutrient conditions.

## Author contribution

MCU: Data curation, Investigation, Methodology, Writing – original draft, Writing – review & editing. HF: Data curation, Investigation, Methodology, Writing – original draft, Writing – review & editing. PM: Data curation, Methodology, Writing – original draft. JVS: Methodology, Writing – original draft. SGE: Data curation, Methodology, AH: Funding acquisition, Supervision, Writing – review & editing.

## Conflict of interest

The authors declare that there is no conflict of interest.

## Funding

This research was supported by grants from Fundação de Amparo a Pesquisa do Estado do Rio de Janeiro (FAPERJ) and Conselho Nacional de Desenvolvimento Científico e Tecnológico (CNPq). CNPq, FAPERJ, and the Coordination for the Improvement of Higher Education Personnel (CAPES) also supported the work with graduate and post-graduate scholarships. MCU and HF were supported by a CNPq postdoctoral fellowship. PM were supported by CNPq/SEMPI/MCTI grant no. 021/2021 (Rhae Program). SGE was supported by a CAPES MSc fellowship. JVS was supported by a CNPq undergraduate fellowship. ASH was supported by a research scientist fellowship from CNPq (306643/2022-7).

## Notes

### Competing Interest Statement

The authors have declared no competing interest.

### Summary of Updates

The data and figures remain unchanged. However, we changed the Title, added a model as Figure 10 and revised the text to improve clarity and overall presentation.

