## Supplemental Figures and Tables for "Silencing ABAP1 INTERACTING PROTEIN 10 (AIP10) enhances root colonization by beneficial bacteria and improves plant performance under nutrient-limited conditions in *Arabidopsis thaliana*"

**Table S1.** Primers used in this study.

| Primer name | Primer Forward Sequence | Primer Reverse Sequence | Organism |
|---|---|---|---|
| **RT-qPCR** | | | |
| *GAPDH* | TTGGTGACAACAGGTCAAGCA | AAACTTGTCGCTCAATGCAATC | *A. thaliana* |
| *UBI14* | TCACTGGAAAGACCATTACTCTTGAA | AGCTGTTTTCCAGCGAAGATG | *A. thaliana* |
| *ABAP1* | TCAGCCTTAAGAAGAGCTTGCA | ACCATAATTGAGAGCTGAGCTTAGTG | *A. thaliana* |
| *Cdt1a* | AATCCGATCACGTCTTGAAGAAG | GAACCACGATCTCAAGAAAGCA | *A. thaliana* |
| *Cyclin B1;1* | CCTCCATTCACTCTCAACAG | CCTGGCAGCTGTGGAATATG | *A. thaliana* |
| *ILA* | CCCGACGGTTATCCAAGTTTC | ACATGTTTCCTCCTACCCACTTACA | *A. thaliana* |
| *LRX4* | CACTCCTCACGCAGATGTATCG | CTGACTTCGGTGTCGGAGAGA | *A. thaliana* |
| *RSL4* | TGGGATGGACATTGGTCTCA | CGTCCGAAGGCGGAGAATA | *A. thaliana* |
| *XHT14* | GCCTCCCCAAGGAGTGTAAAC | CAAGATGAAACAAAACGGAAAGAA | *A. thaliana* |
| *nifH* | AACAAGGCGCAGGAAATCTA | AAATGCCCTTGGAGATGTTG | *H. seropedicae* |
| *nifH* | GCGTCCGCCAGATCGTATT | CTGGGTGGACTGATCTGCAA | *G. diazotrophicus* |
| **q-PCR** | | | |
| *AZO-2* | GCGCGGAAGTCCTGAAT | GGTGGTCGATAACTCCATCG | *A. brasiliense* |
| *FIC* | CGCAAGGCAAACTCCAGATG | CAGGCACACGCATTCTCTTTC | *H. seropedicae* |
| *GSH* | CCTGGCTCGTCTACCTGATC | TGAGGCGGTGACTGACAA | *G. diazotrophicus* |

**Table S2.** Relative abundance of diazotrophic, plant growth–promoting (PGPB), and pathogenic bacterial communities associated with the roots of Col-0 and *aip10-1* plants.

| **Relative Abundance** | | | | |
|---|---|---|---|---|
| ***Diazotrophic bacteria*** | | | | |
| **Organism** | **Root_Col-0** | **oot_*aip10-*** | **Soil_Col-0** | **Soil _*aip10-1*** |
| *Azospirillum brasilense* | 0,00033121 | 0,0004 | 0,00029626 | 0,0002799 |
| *Azospirillum formosense* | 0,00011786 | 0,00013 | 7,3233E-05 | 0,0001954 |
| *Azospirillum halopraeferens* | 0 | 9,1E-05 | 0 | 0 |
| *Azospirillum thiophilum* | 0,0004003 | 0,0004 | 0,0001964 | 0,00022885 |
| *Bradyrhizobium elkanii* | 1,016E-05 | 0,00011 | 3,9945E-05 | 1,0562E-05 |
| *Bradyrhizobium guangxiense* | 9,7535E-05 | 0,00014 | 0,00013981 | 7,0415E-05 |
| *Bradyrhizobium japonicum* | 0,00048158 | 0,00051 | 0,0006824 | 0,00058092 |
| *Burkholderia alpina* | 0,00490522 | 0,00467 | 0,00638126 | 0,00607154 |
| *Burkholderia humi* | 0,0004064 | 0,0004 | 0,0002197 | 0,00031159 |
| *Burkholderia ultramafica* | 8,9407E-05 | 6,6E-05 | 5,6589E-05 | 0,0001021 |
| *Enterobacter tabaci* | 7,9248E-05 | 6,6E-05 | 2,3301E-05 | 7,0415E-06 |
| *Gluconacetobacter johannae* | 0,00012802 | 0,00014 | 0,00042275 | 0,00019188 |
| *Mesorhizobium composti* | 0,00388922 | 0,00364 | 0,00366165 | 0,00442735 |
| *Mesorhizobium denitrificans* | 7,112E-05 | 8,9E-05 | 0,00010652 | 0,00010562 |
| *Mesorhizobium gobiense* | 0,00782722 | 0,00802 | 0,00798573 | 0,00993029 |
| *Mesorhizobium hankyongi* | 5,2832E-05 | 0,00011 | 7,9891E-05 | 0,00010914 |
| *Mesorhizobium robiniae* | 5,8928E-05 | 7,8E-05 | 0,00010319 | 9,8581E-05 |
| *Mesorhizobium sediminum* | 0,00019304 | 0,00022 | 0,00033621 | 0,00033447 |
| *Mesorhizobium silamurunense* | 0,00037795 | 0,00043 | 0,00060251 | 0,00062669 |
| *Microvirga flavescens* | 2,2352E-05 | 6,8E-05 | 7,3233E-05 | 1,2323E-05 |

| ***Z-score*** | | | | |
|---|---|---|---|---|
| ***Diazotrophic bacteria*** | | | | |
| **Organism** | **Root_Col-0** | **Root_aip** | **Soil_Col-0** | **Soil _aip10-1** |
| *Azospirillum brasilense* | 0,1 | 1,4 | -0,6 | -0,9 |
| *Azospirillum formosense* | -0,2 | 0,0 | -1,1 | 1,5 |
| *Azospirillum halopraeferens* | -0,5 | 1,5 | -0,5 | -0,2 |
| *Azospirillum thiophilum* | 0,9 | 0,8 | -1,0 | -1,6 |
| *Bradyrhizobium elkanii* | -0,7 | 1,4 | -0,1 | -0,9 |
| *Bradyrhizobium guangxiense* | -0,4 | 0,9 | 0,8 | -0,5 |
| *Bradyrhizobium japonicum* | -0,9 | -0,6 | 1,3 | 0,0 |
| *Burkholderia alpina* | -0,7 | -1,0 | 1,0 | 6,3 |
| *Burkholderia humi* | 0,8 | 0,8 | -1,3 | -1,1 |
| *Burkholderia ultramafica* | 0,5 | -0,6 | -1,0 | 0,7 |
| *Enterobacter tabaci* | 1,0 | 0,6 | -0,6 | -0,3 |
| *Gluconacetobacter johannae* | -0,7 | -0,6 | 1,5 | -0,1 |
| *Mesorhizobium composti* | 0,0 | -0,7 | -0,7 | 31,2 |
| *Mesorhizobium denitrificans* | -1,3 | -0,3 | 0,8 | 0,0 |
| *Mesorhizobium gobiense* | -0,6 | -0,4 | -0,5 | 52,7 |
| *Mesorhizobium hankyongi* | -1,3 | 0,9 | -0,3 | 1,0 |
| *Mesorhizobium robiniae* | -1,3 | -0,3 | 0,9 | 0,2 |
| *Mesorhizobium sediminum* | -1,1 | -0,6 | 0,9 | 0,5 |
| *Mesorhizobium silamurunense* | -1,1 | -0,6 | 0,8 | 3,7 |
| *Microvirga flavescens* | -0,7 | 0,8 | 0,9 | -1,0 |

| ***PGPB*** | | | | |
|---|---|---|---|---|
| **Organism** | **Root_Col-0** | **oot_*aip10-*** | **Soil_Col-0** | **Soil_*aip10-1*** |
| *Azospirillum brasilense* | 0,00033121 | 0,0004 | 0,00029626 | 0,0002799 |
| *Azospirillum formosense* | 0,00011786 | 0,00021 | 7,3233E-05 | 0,0001954 |
| *Azospirillum thiophilum* | 0,0004003 | 0,0004 | 0,0001964 | 0,00022885 |
| *Bradyrhizobium canariense* | 0,00418793 | 0,00534 | 0,00301254 | 0,00428476 |
| *Bradyrhizobium japonicum* | 0,00048158 | 0,00051 | 0,0006824 | 0,00058092 |
| *Burkholderia alpina* | 0,00467447 | 0,00391 | 0,00638126 | 0,00607154 |
| *Burkholderia humi* | 0,0004064 | 0,0004 | 0,0002197 | 0,00031159 |
| *Erwinia sp.* | 0,00016053 | 0,00011 | 4,3274E-05 | 1,2323E-05 |
| *Paenibacillus medicaginis* | 0,0001016 | 0,00017 | 0,0001731 | 8,8019E-05 |
| *Paenibacillus panacisoli* | 0,00012395 | 0,00013 | 0,00025299 | 6,5134E-05 |
| *Paenibacillus periandrae* | 0,00010973 | 0,00015 | 0,00028295 | 5,2811E-05 |
| *Paenibacillus rigui* | 0,00018085 | 0,00011 | 0,00038947 | 4,2249E-05 |
| *Paraburkholderia diazotrophica* | 0,00047752 | 0,0004 | 0,00035618 | 0,00051931 |
| *Paraburkholderia megapolitana* | 0,00097942 | 0,00088 | 0,00052927 | 0,00137662 |
| *Paraburkholderia metrosideri* | 0,00412493 | 0,00386 | 0,00436069 | 0,00444319 |
| *Paraburkholderia oxyphila* | 4,2672E-05 | 0,00146 | 0,00658764 | 0,0025191 |
| *Paraburkholderia pallidirosea* | 0,00060756 | 0,00058 | 0,00061249 | 0,00052459 |
| *Paraburkholderia xenovorans* | 6,2992E-05 | 0,00011 | 0,00036617 | 0,00012323 |
| *Pseudomonas sp.* | 0,00017881 | 0,00011 | 0,00013981 | 8,0977E-05 |

| ***PGPB*** | | | | |
|---|---|---|---|---|
| **Organism** | **Root_Col-0** | **Root_aip** | **Soil_Col-0** | **Soil_aip10-1** |
| *Azospirillum brasilense* | 0,1 | 1,4 | -0,6 | -0,9 |
| *Azospirillum formosense* | -0,2 | 0 | -1,1 | 1,3 |
| *Azospirillum thiophilum* | 0,9 | 0,8 | -1 | -0,7 |
| *Bradyrhizobium canariense* | 0,8 | -0,6 | -1,1 | 0,9 |
| *Bradyrhizobium japonicum* | -0,9 | -0,6 | 1,3 | 0,2 |
| *Burkholderia alpina* | -0,7 | -1 | 1 | 0,7 |
| *Burkholderia humi* | 0,8 | 0,8 | -1,3 | -0,3 |
| Erwinia sp. | 1,2 | 0,5 | -0,6 | -1,0 |
| *Paenibacillus medicaginis* | -0,7 | 0,9 | 0,9 | -1 |
| *Paenibacillus panacisoli* | -0,3 | -0,1 | 1,4 | -1 |
| *Paenibacillus periandrae* | -0,4 | 0 | 1,4 | -1 |
| *Paenibacillus rigui* | 0 | -0,5 | 1,4 | -0,9 |
| *Paraburkholderia diazotrophica* | 0,5 | -0,5 | -1,1 | 1,1 |
| *Paraburkholderia megapolitana* | 0,1 | -0,2 | -1,2 | 1,2 |
| *Paraburkholderia metrosideri* | -0,3 | -1,3 | 0,6 | 0,9 |
| *Paraburkholderia oxyphila* | -0,9 | -0,4 | 1,4 | 0 |
| *Paraburkholderia pallidirosea* | 0,7 | -0,1 | 0,8 | -1,4 |
| *Paraburkholderia xenovorans* | -0,7 | -0,4 | 1,5 | -0,3 |
| *Pseudomonas sp.* | 1,2 | -0,4 | 0,3 | -1,1 |

| ***Pathogenic bacteria*** | | | | |
|---|---|---|---|---|
| **Organism** | **Root_Col-0** | **oot_*aip10-*** | **Soil_Col-0** | **Soil _*aip10-1*** |
| *Agrobacterium fabrum* | 0 | 0 | 1,3315E-05 | 1,0562E-05 |
| *Agrobacterium larrymoorei* | 0 | 0 | 4,6603E-05 | 0 |
| *Erwinia billingiae* | 0 | 0 | 8,6548E-05 | 0 |
| *Erwinia soli* | 3,8608E-05 | 0 | 1,6496E-05 | 1,4083E-05 |
| *Erwinia sp.* | 0,00016053 | 0,00011 | 4,3274E-05 | 1,2323E-05 |
| *Erwinia teleogrylli* | 0 | 0 | 2,1125E-05 | 2,1125E-05 |
| *Pantoea brenneri* | 8,2478E-06 | 0 | 0 | 0 |
| *Pseudomonas aeruginosa* | 0 | 0 | 9,9863E-06 | 0 |
| *Pseudomonas sp.* | 0,00017881 | 0,00011 | 0,00013981 | 8,0977E-05 |
| *Xanthomonas arboricola* | 8,128E-06 | 0 | 0 | 0 |
| *Xanthomonas axonopodis* | 1,016E-05 | 0 | 1,6644E-05 | 5,2811E-06 |
| *Xanthomonas campestris* | 0 | 0 | 6,6575E-06 | 0 |
| *Xanthomonas citri* | 1,016E-05 | 1E-05 | 2,663E-05 | 1,9364E-05 |
| *Xanthomonas translucens* | 0 | 0 | 1,3315E-05 | 0 |

| ***Pathogenic bacteria*** | | | | |
|---|---|---|---|---|
| **Organism** | **Root_Col-0** | **Root_aip** | **Soil_Col-0** | **Soil _aip10-1** |
| *Agrobacterium fabrum* | -0,9 | -0,9 | 0,8 | 0,9 |
| *Agrobacterium larrymoorei* | -0,5 | -0,5 | 1,5 | -0,5 |
| *Erwinia billingiae* | -0,5 | -0,5 | 1,5 | -0,5 |
| *Erwinia soli* | 0,4 | -1,5 | 0,5 | 0,6 |
| *Erwinia sp.* | -0,9 | -0,6 | 0,2 | 1,3 |
| *Erwinia teleogrylli* | -0,5 | -0,5 | -0,5 | 1,5 |
| *Pantoea brenneri* | 1,5 | -0,5 | -0,5 | -0,5 |
| *Pseudomonas aeruginosa* | -0,5 | -0,5 | 1,5 | -0,5 |
| *Pseudomonas sp.* | -1,1 | 0,3 | -0,4 | 1,2 |
| *Xanthomonas arboricola* | 1,5 | -0,5 | -0,5 | -0,5 |
| *Xanthomonas axonopodis* | 0,5 | -1,5 | 0,4 | 0,6 |
| *Xanthomonas campestris* | -0,5 | -0,5 | 1,5 | -0,5 |
| *Xanthomonas citri* | 0,8 | 0,8 | -1,2 | -0,5 |
| *Xanthomonas translucens* | -0,5 | -0,5 | 1,5 | -0,5 |

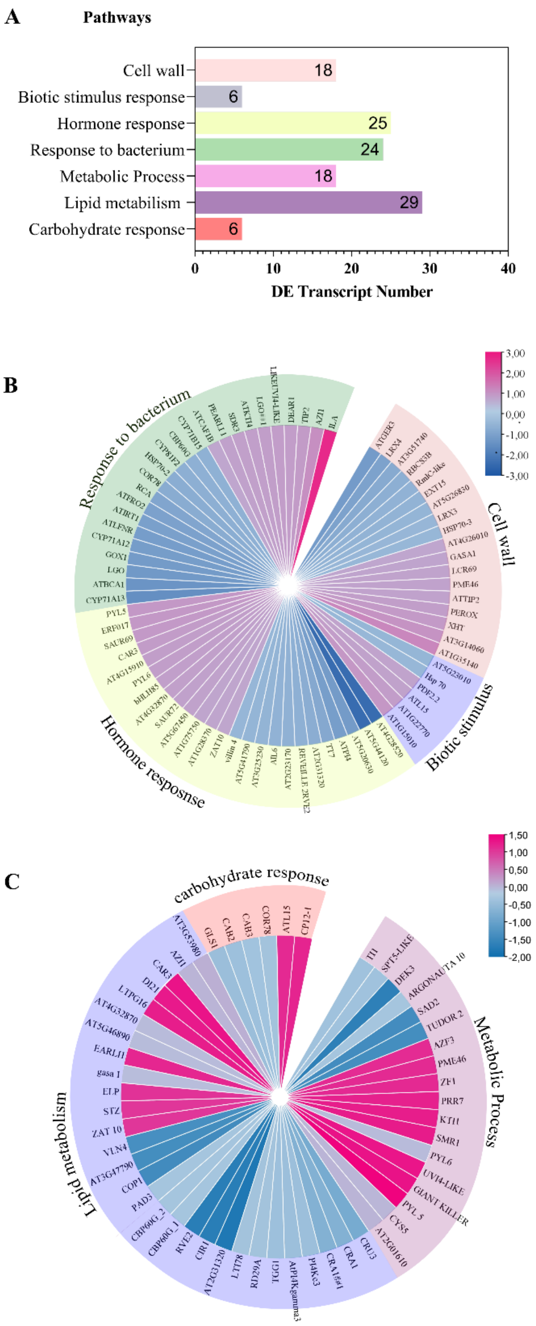


**Fig S1. Differential gene expression patterns in roots Col-0 and *aip10-1* of *A. thaliana,* 11 days after germination**. **(A)** Functional classification of differentially expressed transcripts (DEGs) in metabolic and biological pathways. **(B)** Gene expression pattern within the categories response to bacteria, hormone response, biotic stimulus response, and cell wall, highlighting the contribution of genes related to plant immunity and structural remodeling. **(C)** Expression pattern of genes involved in carbohydrate response, lipid metabolism, and general metabolic processes, suggesting an intense reorganization of energy and structural metabolism during root development. Expression values were obtained from RPKM data and represented as *log fold change* between *aip10-1*vs Col-0 roots. Pink bars indicate higher gene induction in *aip10-1*, while blue bars represent gene repression.

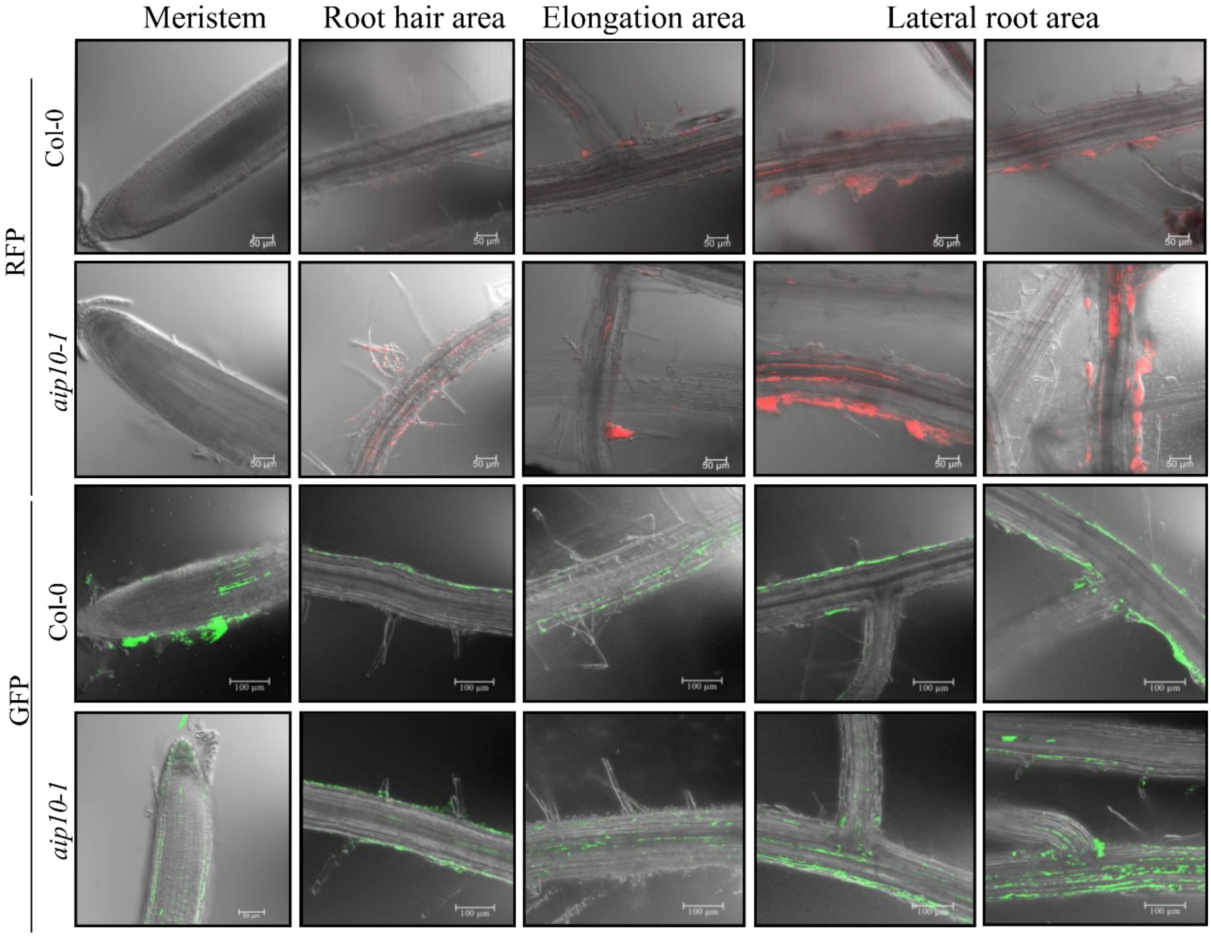


**Fig S2. Visualization of bacterial colonization patterns in different root zones of *A. thaliana* Col-0 and *aip10-1* mutant plants.** Confocal microscopy images showing bacterial localization in the meristematic zone, root hair area, elongation zone, and lateral root area. Plants were inoculated with *G. diazotrophicus* PAL5 strain expressing RFP (red) or *H. seropedicae* HRC54 strain expressing GFP (green) reporters, enabling direct visualization of colonization along the root surface and tissues. In *aip10-1* mutant roots, a higher fluorescence signal was consistently observed across all analyzed regions, indicating a greater bacterial presence and colonization density compared with the Col-0 wild type. Scale bars = 100 µm.

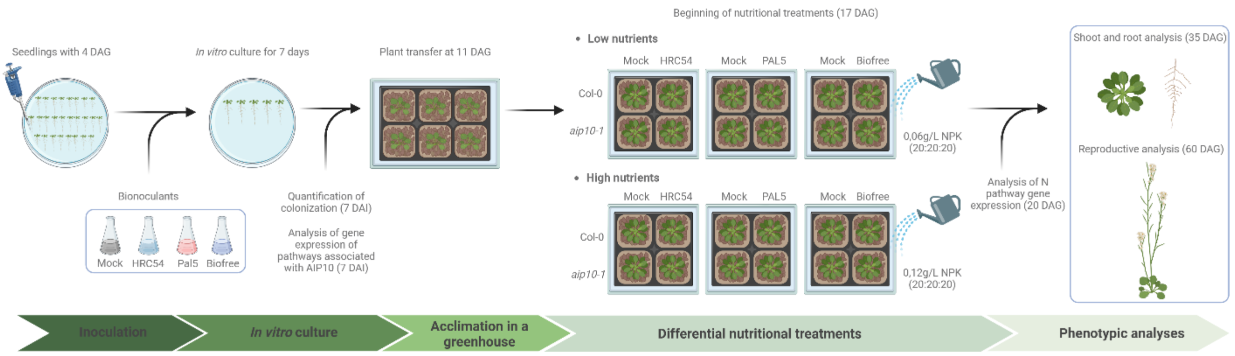


**Fig S3. Experimental design of the differential fertilization trial in *aip10-1* and wild-type Col-0 plants inoculated with *H. seropedicae* HRC54 (HRC54), *G. diazotrophicus* Pal5 (Pal5), Biofree or non-inoculated (mock).** Seeds were pre-germinated *in vitro*, and seedlings were inoculated at 4 DAG. Seven days after inoculation (11 DAG), plants were transferred to the greenhouse. Differential irrigation treatments with low nutrients (0.06 g/L NPK; 20:20:20) and high nutrients (0.12 g/L NPK; 20:20:20) were initiated at 17 DAG. Samples for molecular analyses were collected at 11 and 20 DAG, while those for phenotypic analyses were collected at 35 and 60 DAG. Created with BioRender.com.

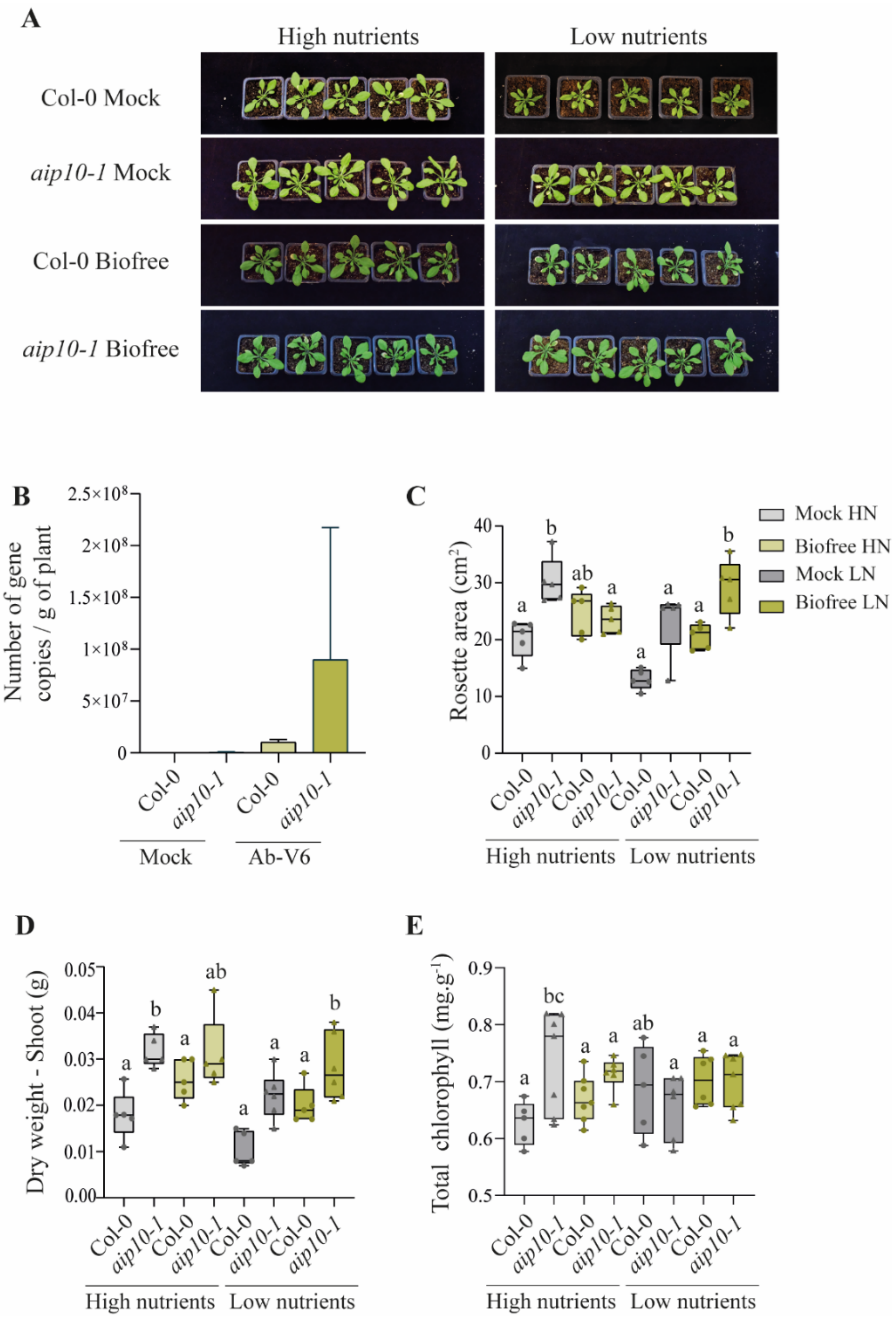


**Fig S4. Growth performance and physiological responses of Col-0 and *aip10-1* plants inoculated with commercial bioinoculant *Biofree* under contrasting nutrient conditions. (A)** Representative images of Col-0 and *aip10-1* plants grown under high-nutrient (HN) and low-nutrient (LN) conditions, and non-inoculated (Mock) or inoculated with the *Biofree* bacterial consortium. **(B)** Absolute qPCR quantification of *Azospirillum brasilense* Ab-V6 colonization in roots of Col-0 and aip10-1 at 7 DAI. **(C–E)** Quantification of key growth traits, including rosette area **(C)**, shoot dry weight **(B)**, and total chlorophyll content **(E)**. Plants were evaluated at the vegetative stage, and boxplots depict the distribution of five biological replicates. Statistical analysis was performed using the Student's t-test (P ≤ 0.05). Different letters indicate statistically significant differences ($P \leq 0.05$).

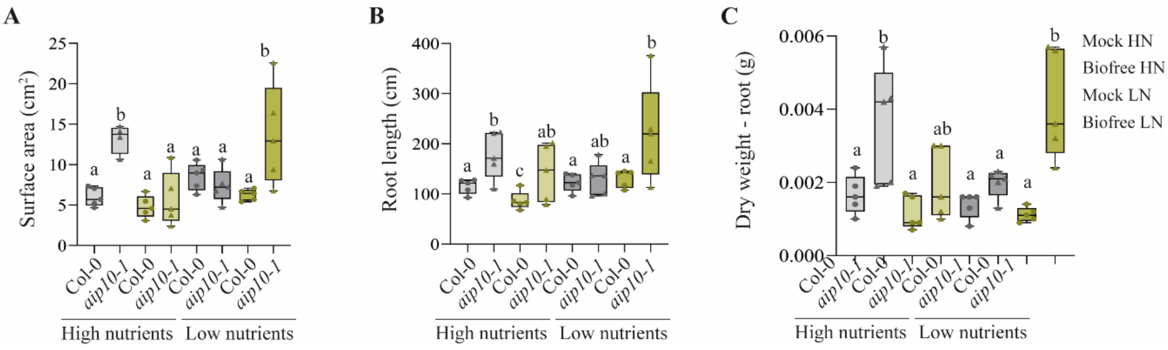


**Fig S5. Root architecture responses of Col-0 and *aip10-1 A. thaliana* plants to *Biofree* inoculation under contrasting nutrient conditions.** Quantitative analysis of **(A)** root dry weight, **(C)** primary root length and **(D)** total root surface area. Boxplots represent five biological replicates; statistical analysis was performed using the Student's t-test (P ≤ 0.05); different letters indicate statistically significant differences ($P \leq 0.05$).

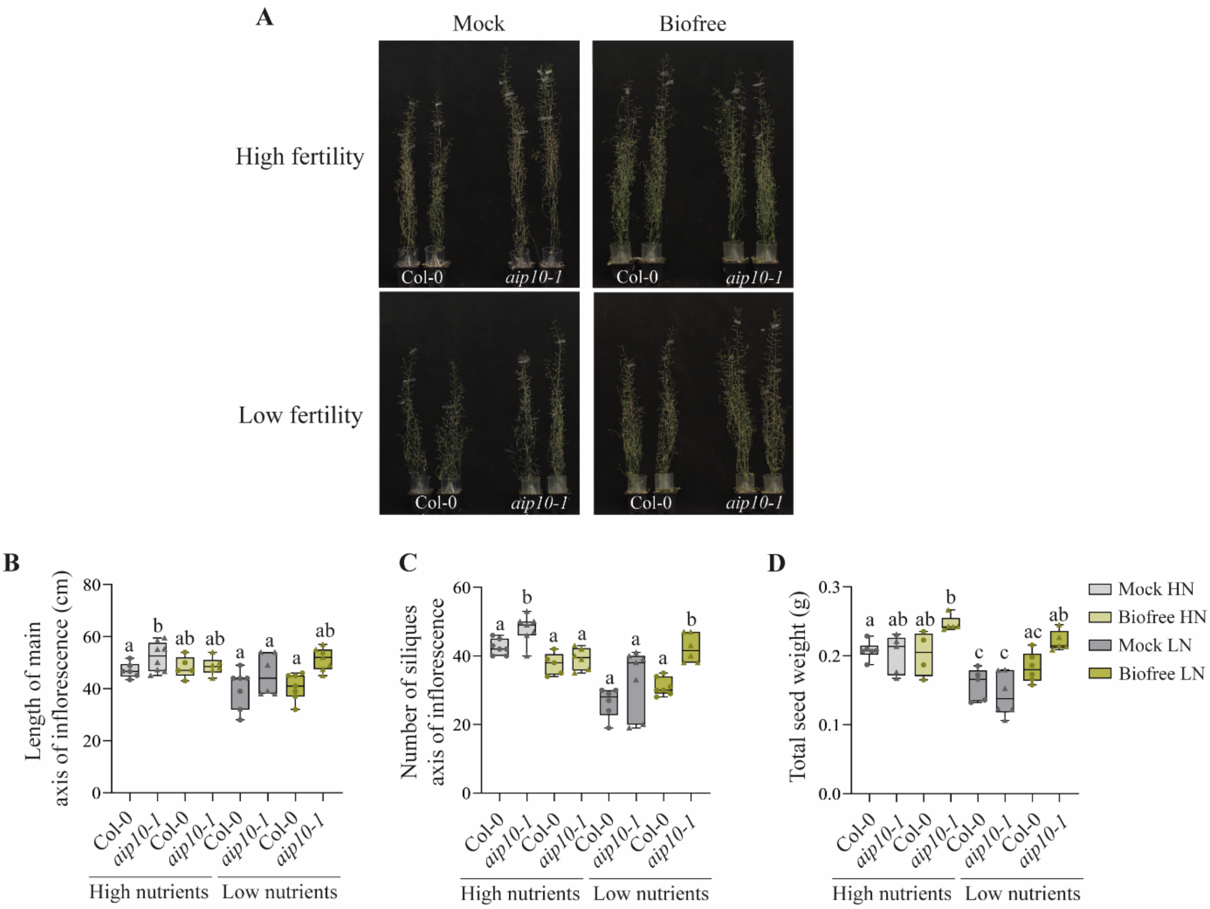


**Fig S6. Reproductive performance of Col-0 and *aip10-1 A. thaliana* plants in response to *Biofree* inoculation under high- and low-nutrient conditions. (A)** Representative images of inflorescence Col-0 and *aip10-1* grown under high-fertility and low-fertility conditions, non-inoculated (Mock) or inoculated with the *Biofree* microbial consortium. **(B–D)** Quantification of reproductive traits including. **(B)** main inflorescence length. **(C)** number of siliques per main inflorescence, and **(D)** total seed weight per plant. Data are presented as boxplots of five biological replicates. Statistical analysis was performed using the Student's t-test ($P \leq 0.05$). Different letters denote significant differences ($P \leq 0.05$).
